# Impaired function and delayed regeneration of dendritic cells in COVID-19

**DOI:** 10.1101/2021.05.26.445809

**Authors:** Elena Winheim, Linus Rinke, Konstantin Lutz, Anna Reischer, Alexandra Leutbecher, Lina Wolfram, Lisa Rausch, Jan Kranich, Paul R. Wratil, Johanna E. Huber, Dirk Baumjohann, Simon Rothenfußer, Johannes C. Hellmuth, Clemens Scherer, Maximilian Muenchhoff, Michael von Bergwelt-Baildon, Konstantin Stark, Tobias Straub, Thomas Brocker, Oliver T. Keppler, Marion Subklewe, Anne B. Krug

## Abstract

Disease manifestations in COVID-19 range from mild to severe illness associated with a dysregulated innate immune response. Alterations in function and regeneration of dendritic cells (DC) and monocytes may contribute to immunopathology and influence adaptive immune responses in COVID-19 patients. We analyzed circulating DC and monocyte subsets in 65 hospitalized COVID-19 patients with mild/moderate or severe disease from acute disease to recovery and in healthy controls. Persisting reduction of all DC subpopulations was accompanied by an expansion of proliferating Lineage^-^ HLADR^+^ cells lacking DC markers. Increased frequency of the recently discovered CD163^+^ CD14^+^ DC3 subpopulation in patients with more severe disease was associated with systemic inflammation, activated T follicular helper cells, and antibody-secreting cells. Persistent downregulation of CD86 and upregulation of PD-L1 in conventional DC (cDC2 and DC3) and classical monocytes associated with a reduced capacity to stimulate naïve CD4^+^ T cells correlated with disease severity. Long-lasting depletion and functional impairment of DCs and monocytes may have consequences for susceptibility to secondary infections and therapy of COVID-19 patients.

## Introduction

Coronavirus disease 2019 (COVID-19), caused by novel severe acute respiratory syndrome coronavirus (SARS-CoV-2), has emerged in December 2019 (1) and is currently causing a global health emergency. COVID-19 is characterized by diverse clinical manifestations ranging from asymptomatic, mild, moderate, to severe disease, including pneumonia which may progress to acute respiratory distress syndrome and multi-organ failure (2). Exacerbated systemic inflammatory responses and thrombophilia frequently leading to cardiovascular complications are hallmarks of the severe form of the disease (3). Several contributors to a more severe disease outcome have been identified so far, such as age, male sex, comorbidities, immunosuppression, autoantibodies against type I IFN and genetic variants. The disease course is strongly influenced by the dynamic interaction of the virus with the immune system (4, 5). Disease severity was shown to correlate strongly with reduced lymphocyte and increased neutrophil counts in the blood as well as high concentrations of inflammatory cytokines such as IL-6, TNF-*α*, IL-1*β* and chemokines (e.g. CXCL10 and CCL2) (2, 6, 7). Antibody and T cell responses were found in over 90 % of convalescent individuals (5, 8–10) including T follicular helper cell activation and plasma cell expansion (10–12). Immunological memory develops after natural infection lasting at least 6-8 months (13, 14).

As highly efficient antigen-presenting cells (APCs), DCs are essential in recognizing pathogens, orchestrating innate and adaptive immune responses and secreting inflammatory mediators. Each DC subpopulations has specific functions in the antiviral immune response. Conventional DC (cDC) are highly efficient in presenting antigens and stimulating naïve T cells to expand and differentiate. While cDC1 are specially equipped for cross-presentation of antigens to CD8^+^ T cells, cDC2 shape Th cell responses (15). DC3 in human blood share characteristics of both cDC2 and monocytes but are distinct in ontogeny and may exert specific functions, but their roles in peripheral tissues and during immune responses are still unclear. In COVID-19 patients, an overall reduction of cDC subsets in the blood was observed in several studies (16–19) and activated cDC2 were found to accumulate in the lungs of critically ill COVID-19 patients (18). Plasmacytoid DC (pDC), which rapidly produce antiviral type I interferons and inflammatory chemokines are reduced in numbers and functionally impaired in COVID-19 patients (7, 16, 17, 20, 21). Monocytes are quickly recruited to inflammation sites and can differentiate into macrophages and monocyte-derived DCs (22). In COVID-19, the recruitment of monocytes into the inflamed lung and subsequent production of proinflammatory cytokines could contribute to disease progression and tissue damage (18, 23–25). However, in patients with severe COVID-19 monocytes and DCs in the blood were found to express lower levels of HLADR and CD86 (16, 17, 19, 20, 26–30).

In this study, we sought to gain an in-depth understanding of the dynamic changes in frequencies, activation status, and functionality of blood monocyte and DC subsets in correlation with adaptive immune responses and disease severity in COVID-19 patients. We observed a long-lasting reduction of DC subpopulations with an expansion of proliferating Lineage^-^ HLADR^+^ cells lacking DC markers and delayed regeneration. High-dimensional longitudinal flow cytometric analysis revealed an early type I IFN induced response and a longer-lasting PD-L1^hi^ CD86^lo^ phenotype in DC3 and classical monocytes. This dysregulated activation was associated with a reduced ability to stimulate T cells and correlated with disease severity. CD163^+^ CD14^+^ cells within DC3 increased in the patients with more severe disease and correlated with inflammation and subsequent activation of Tfh cells and B cells, but not antibody titers. Our results provide evidence for long-lasting aberrant activation and delayed regeneration of circulating APCs in COVID-19.

## Results

### Persisting reduction of circulating DC subpopulations in COVID-19 patients

From 65 patients with PCR-confirmed SARS-CoV-2 infection, a total of 124 samples of PBMC were used for flow cytometric analysis. Patients with active COVID-19 (mild/moderate or severe) were compared with recovered patients and a control group including healthy donors and SARS-CoV-2-negative patients (see Fig. 1a and Table S1 for a detailed description of the cohorts). COVID-19 severity was assessed using an ordinal scale from 1 to 8 adopted from the World Health Organization (31). The maximal value (WHO max) reached by the patients in our cohort correlated with laboratory markers of inflammation and altered peripheral blood leucocyte composition that are associated with disease severity (Fig. 1b). We first characterized monocytes and DCs in PBMCs by multi-dimensional flow cytometry (Fig. 2a). In line with published observations, we observed a relative reduction of monocytes in patients with mild/moderate disease and an increase of low-density neutrophils within the PBMC in a subgroup with more severe disease (Figure 2b, S1) (28). The percentage of cells within the DC gate (Lin^-^ HLADR^+^ CD14^-^ CD88/89^-^ CD16^-^) tended to be lower in patients than in controls. Within CD88/89^+^ monocytes we found a relative increase of classical CD14^+^ CD16^-^ monocytes (mo 1) and a decrease of CD14^lo^ CD16^+^ non-classical monocytes (mo 2) in patients with mild/moderate and severe disease in our cohort (Fig. S4), confirming published results (28, 29). Mo 2 were significantly reduced and mo 1 concomitantly increased compared to controls within the first 15 days after diagnosis recovering thereafter (Fig. S4).

**Fig. 1.**
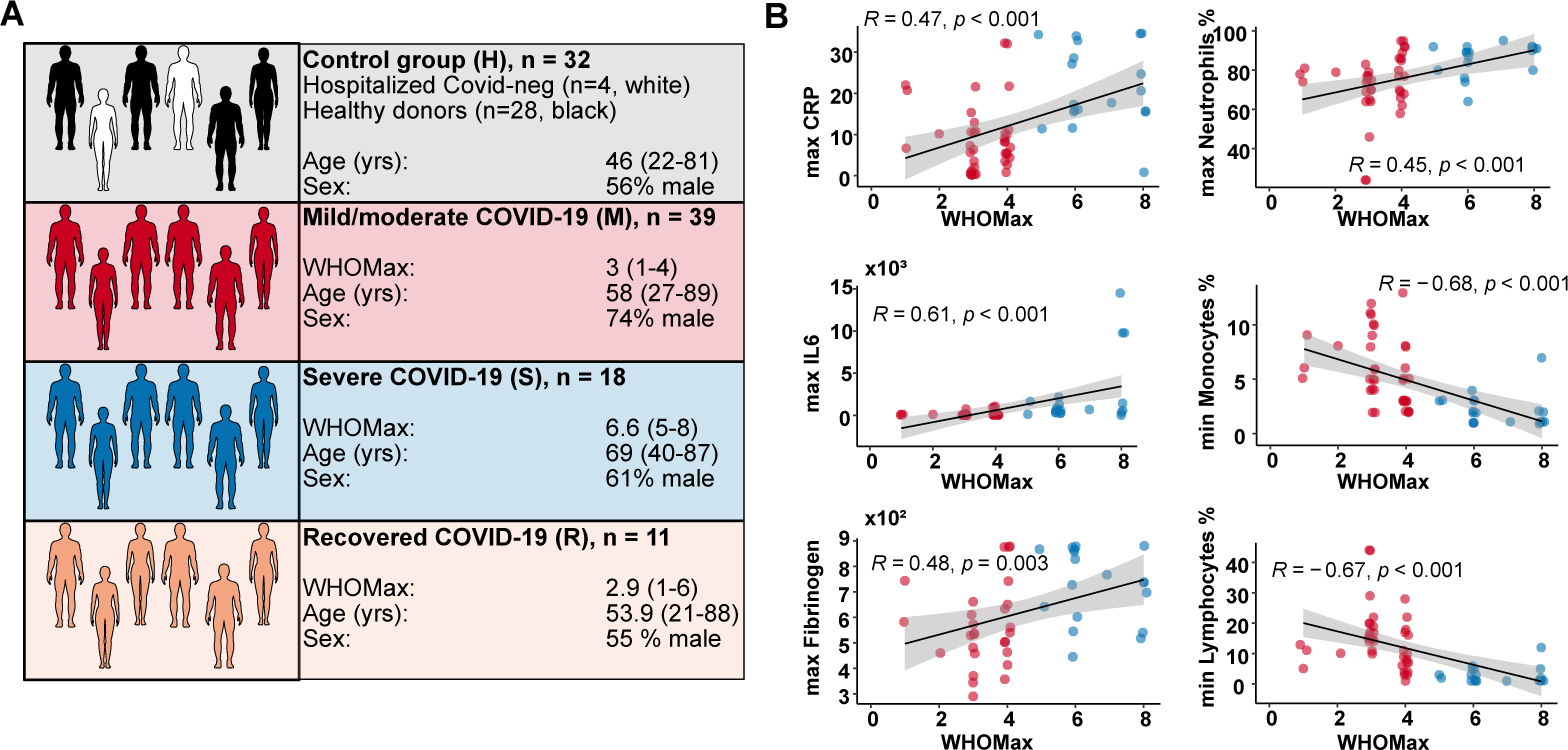
Characterization of the study cohorts. (A) The number, age, sex and maximal WHO ordinal scale (WHO max) reached are shown for the four different study groups. The control group (H) contained 28 healthy blood donors (black) and 4 SARS-CoV-2-negative patients (white). Patients with acute COVID-19 were grouped into mild/moderate (M, red, n=39) and severe (S, blue, n=18). A group of recovered patients was included for comparison (R, orange, n=11). (B) Correlation analysis of WHO max values with routine laboratory values (minimal and maximal values reached during hospitalization). CRP, C-reactive protein. Spearman’s rank correlation coefficients, p-values and linear regression lines are shown.

**Fig. 2.**
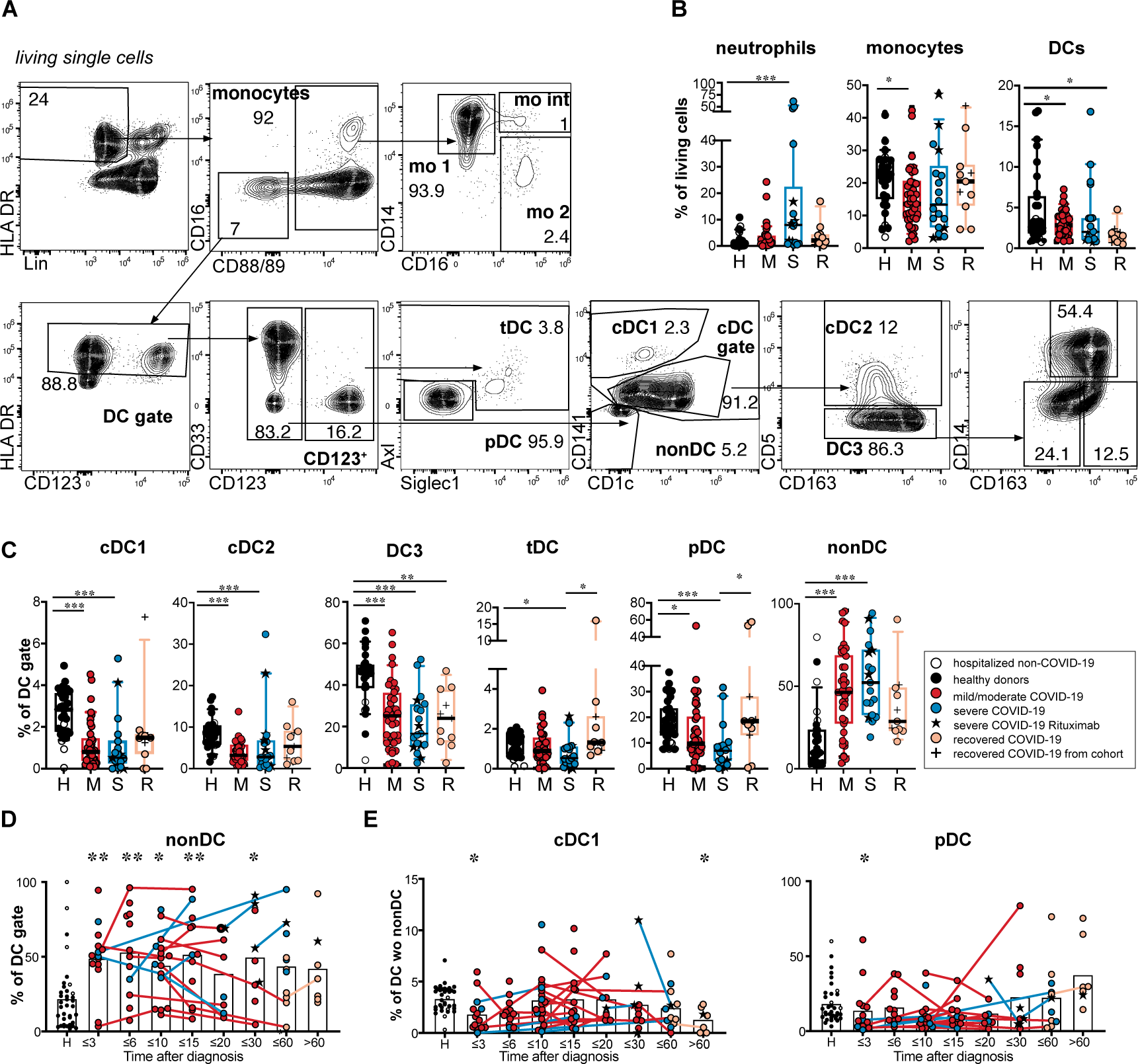
Reduction of DC subpopulation and expansion of immature HLADR^+^ cells in COVID-19 patients. (A) Gating strategy for DC and monocyte subtypes in the blood: Within HLADR^+^ Lineage (CD3, CD15, CD19, CD20, CD56, CD66b), negative (Lin^-^) cells monocytes were gated as CD88/89 positive cells and separated into mo 1 (CD14^+^ CD16^-^ classical monocytes, mo int (CD14^+^ CD16^+^ intermediate monocytes, mo 2 (CD14^lo^ CD16^+^ non-classical monocytes). HLADR^+^ Lin^-^ CD88/89^-^ CD16^-^ cells were regated on HLADR positive cells (DC gate). Within the CD123^+^ DC fraction pDCs (Siglec1^-^ Axl^-^) and tDCs (Axl^+^ and/or Siglec1^+^) were distinguished. Within the CD123^-^ DC fraction cDC1 (CD141+ CD1c^lo^), cDC2 (CD141^-^, CD1c^+^, CD5^+^), DC3 (CD141^-^CD1c+ CD5^-^ DCs) and “non-DC” (CD141^-^ CD1c^-^) were identified. DC3 were further separated into CD163^-^ CD14^-^, CD163^+^ CD14^-^ and CD163^+^ CD14^+^ DC3 subsets. (B) Percentage of neutrophils (Lin^+^ CD88/89^+^ CD16^+^), monocytes (Lin^-^, HLADR^+^, CD88/CD89^+^) and DCs (Lin^-^, HLADR^+^, CD88/CD89^-^) of living PBMC. Healthy donors (=H, black symbols, n=28), hospitalized COVID-19 negative patients (=white symbols, n=4), acute COVID-19 patients with mild/moderate (=M, red symbols, n=39), severe (=S, blue symbols, n=18) disease at the first analysis timepoint and recovered patients (orange, n=11). In the severe group, patients that had received B cell-depleting therapy before diagnosis (n=5) are marked by a black star. Recovered patients that had already been analyzed during acute disease and were sampled again after recovery are indicated by a plus sign. (C) Relative frequencies of DC subsets and non-DCs within the DC gate are shown (Kruskal-Wallis test with Dunn’s correction, n=97-100). (D) Frequency of non-DCs within the DC gate at different grouped time points after diagnosis. (E) Frequency of cDC1 and pDCs within the differentiated DC population (after excluding CD141^-^ CD1c^-^ non-DCs) at different grouped time points after diagnosis. (D and E) Connected lines represent multiple measurements of the same patient at different time points. Columns indicate the mean. Colors and symbols as in C. Comparison of the indicated time points with the healthy control group (Kruskal-Wallis test with Dunn’s correction, n=124-127). Statistical significance in B, C, D, E is indicated by * p< 0.05, ** p> 0.01, *** p<0.001.

Focusing on DCs, we found a significant relative reduction of cDC1, cDC2, and pDC within the Lin^-^ HLADR^+^ CD14^-^ CD88/89^-^ CD16^-^ population in COVID-19 patients compared to controls. tDCs showed significantly lower frequency in severe patients (Fig. 2 c). At the same time, a population of cells lacking typical DC markers such as CD1c, CD141, CD123, and CD11c but expressing HLADR and partially CD86 (Fig. 2c and Fig. S1), after that called non-DCs, was found to be significantly expanded within the Lin^-^ HLADR^+^ CD14^-^ CD88/89^-^ CD16^-^ fraction. Uniform Manifold Approximation and Projection (UMAP) analysis showed that these cells cluster separately from differentiated DC subpopulations (Fig. S1). We analyzed the expression of several markers of known progenitor cells and found this population to be CD34^-^ CD127^-^ CD117^-^ CD115^-^ with varying expression of CD45RA and detection of proliferation marker Ki67 (Fig. S1). This proliferative HLADR^+^ CD86^+/–^ population, therefore, did not phenotypically overlap with a defined progenitor population. The increased frequency of this DC-like population was long-lasting and could still be observed even in recovered patients more than 60 days after primary diagnosis of COVID-19 (Fig. 2d).

Changes in blood DC numbers and subset composition after bacterial or viral infection are highly dynamic (32). We, therefore, analyzed the frequencies of DC subsets within total DCs (excluding the non-DC fraction) at different time points (Fig. 2e). While the percentages of cDC2, tDC and DC3 within differentiated DCs (after exclusion of non-DCs) were not consistently altered in patients versus controls, the frequencies of cDC1 and pDCs were significantly reduced at the earliest time points (*≤* 3 days after diagnosis) and largely restored in recovered patients (Fig. 2e). Our results show that all subpopulations of differentiated circulating DCs are relatively reduced with cDC1 and pDCs being most affected.

### Shift towards CD163^+^ CD14^+^ cells within DC3 correlates with COVID-19 disease activity and inflammatory markers

DC3, which share phenotypic and functional features of cDC2 and monocytes, represent the largest subpopulation of DCs in the blood. Differential expression of CD163 and CD14 marks different stages of maturation and activation in DC3 (33, 34), and an increased frequency of CD163^+^ CD14^+^ blood DC3 with proinflammatory function exists in patients with active SLE. We, therefore, hypothesized that the CD163^+^ CD14^+^ fraction within DC3 is expanded also in COVID-19 patients. We observed a significantly increased frequency of CD163^+^ CD14^+^ cells and decreased frequency of CD163^+^ CD14^-^ cells in the DC3 subset in COVID-19 patients compared to controls. This shift was most pronounced in patients with severe disease (Fig. 3a-d) and in samples taken up to 20 days after diagnosis (Fig. 3c). The percentage of CD163^+^ CD14^+^ cells within DC3 returned to the level of healthy controls in the majority of recovered patients (Fig. 3b and c). The frequency of CD163^+^ CD14^+^ DC3 correlated positively, and the frequency of CD163^+^ CD14^-^ DC3 correlated negatively with disease severity (WHO Max), maximal CRP, and maximal IL-6 values during hospitalization and with the actual CRP values at the time of sampling (Fig. 3e 6 and f). Thus, the shift towards more mature CD163^+^ CD14^+^ DC3 in COVID-19 patients was a persistent phenotype associated with inflammation and higher disease activity.

**Fig. 3.**
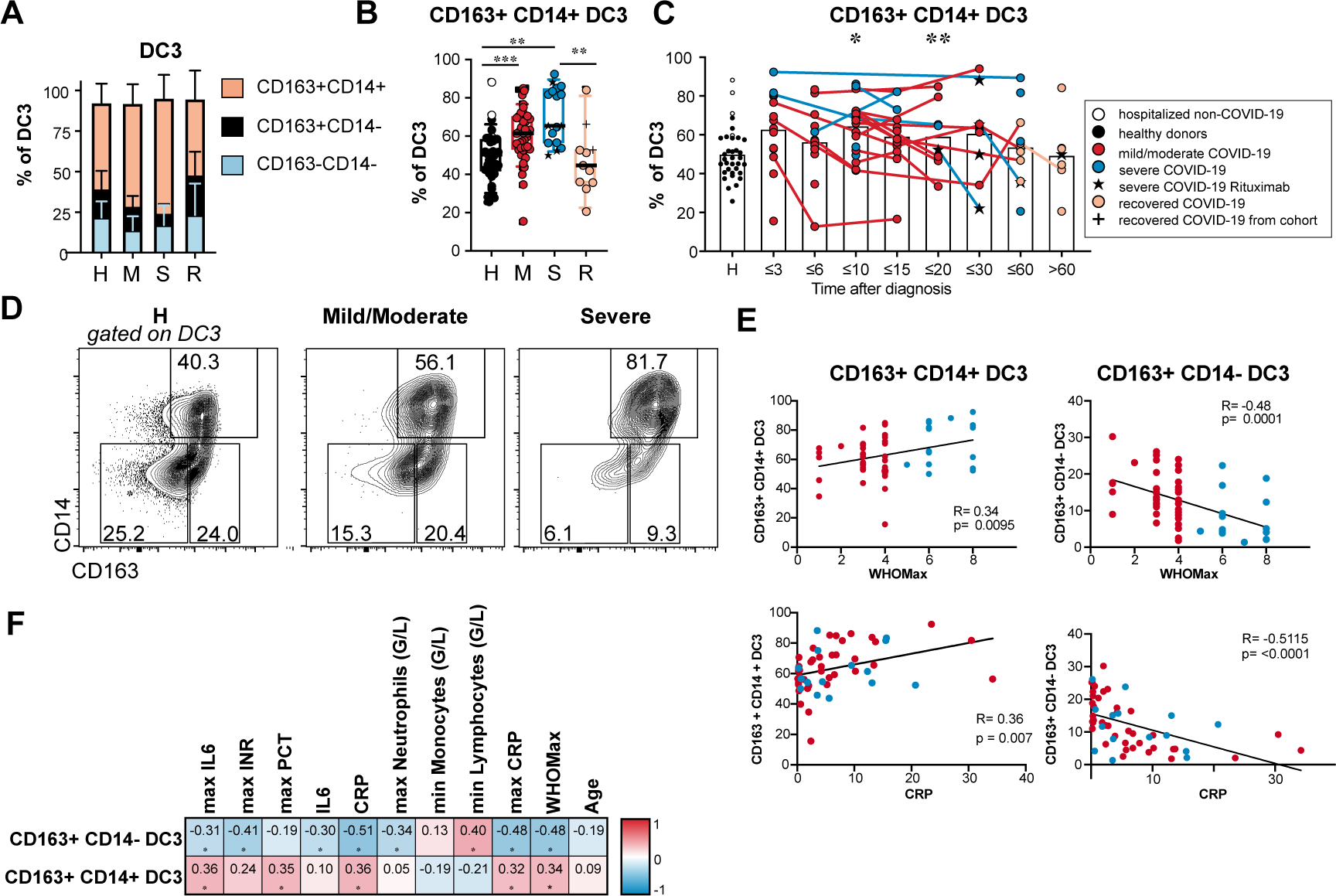
Increased percentage of CD163^+^ CD14^+^ DC3 in COVID-19 patients. (A, B) Frequencies of DC3 subtypes identified by CD163 and CD14 expression are shown for healthy/non-COVID donors (H, n=31), patients with mild/moderate (M, n=39) and severe disease (S, n=18) at the first analysis timepoint and recovered patients (R, n=11). (B) Results for individual patients are indicated by symbols as in Fig. 2 (Kruskal-Wallis test with Dunn’s correction, n=99). (C) Frequency of CD163^+^ CD14^+^ cells within DC3 in all patients of the cohort at different grouped time points after diagnosis. Connected lines represent multiple measurements of the same donor at different time points. Columns indicate the mean (Kruskal-Wallis test with Dunn’s correction, n=124). *p<0.05, ** p> 0.01, *** p<0.001. (D) CD14 and CD164 expression in DC3. Representative results of one healthy donor and two patients with moderate and severe COVID-19 are shown. (E) Correlation of relative frequencies of CD163^+^ CD14^+^ and CD163^+^ CD14^-^ DC3 with WHO max score (n=57) and CRP concentration in the plasma (n=55) at the same time point. Spearman’s rank correlation coefficients, p-values and linear regression lines are shown. (F) Correlation with inflammatory markers, blood cell counts, disease severity and age. Spearman correlation coefficients (-1 to 1) and adjusted p-values are shown.

### Early transient expression of Siglec-1 and persistent CD86^lo^ PD-L1^hi^ phenotype of circulating cDCs and monocytes in COVID-19

In addition to the described dynamic changes in DC and monocyte subset composition after SARS-CoV-2 infection, we postulated that the expression of costimulatory molecules, activation markers and chemokine receptors in these cell types is altered in patients with active COVID-19. A high-dimensional spectral flow cytometry analysis was performed on PBMC of 20 patients with mild/moderate disease, 6 patients with severe disease and 11 healthy donors (see Table S1 for a description of this subcohort). Expression levels of the indicated markers were compared between these 3 groups for each DC and monocyte subpopulation (shown in the heatmap in Fig. 4a). Costimulatory molecule CD86 was downregulated in cDC subsets, mo 1 and mo int populations in patients compared to controls. HLADR expression in mo 1 and DC3 was downregulated only in severe disease and upregulated in mild/moderate disease. At the same time, CD40 and programmed death-ligand 1 (PD-L1) expression in DC and monocyte subsets were increased in both patient groups indicating opposing expression of costimulatory and regulatory molecules (Figure 4a and b). The PD-L1/CD86 ratio in DC3 was increased in patients until late time points (Fig. 4c). It correlated with inflammatory markers and disease severity and segregated patients from controls and patients with mild disease from patients with more severe disease in principal component analysis (Fig. 4h and i). Investigating the whole cohort (65 patients), we detected a distinct CD86^lo^ PD-L1^hi^ DC3 subpopulation, which was also present in cDC2 but not in monocyte populations (Fig. 4b and d). These CD86^lo^ PD-L1^hi^ DC3 and cDC2 were significantly enriched in patients with severe COVID-19 (Fig. 4d). Considering all samples measured, a sizable population of CD86^lo^ PD-L1^hi^ DC3 (> 10%) was found in 7.8 % of samples of the mild/moderate group, 43.3. % of the severe group, 33.3% of the recovered group and 8.3 % of controls (data not shown). This subpopulation had expanded in COVID-19 patients with and without glucocorticoid therapy (Fig. S2). None of the healthy donors in the control group, but 3 non-COVID-19 control patients had more than 10% of the CD86^lo^ PD-L1^hi^ DC3. Two control patients with a high percentage of this subset suffered from COPD and interstitial lung disease indicating that this subset can also be found in other pathologies associated with prolonged inflammatory responses.

**Fig. 4.**
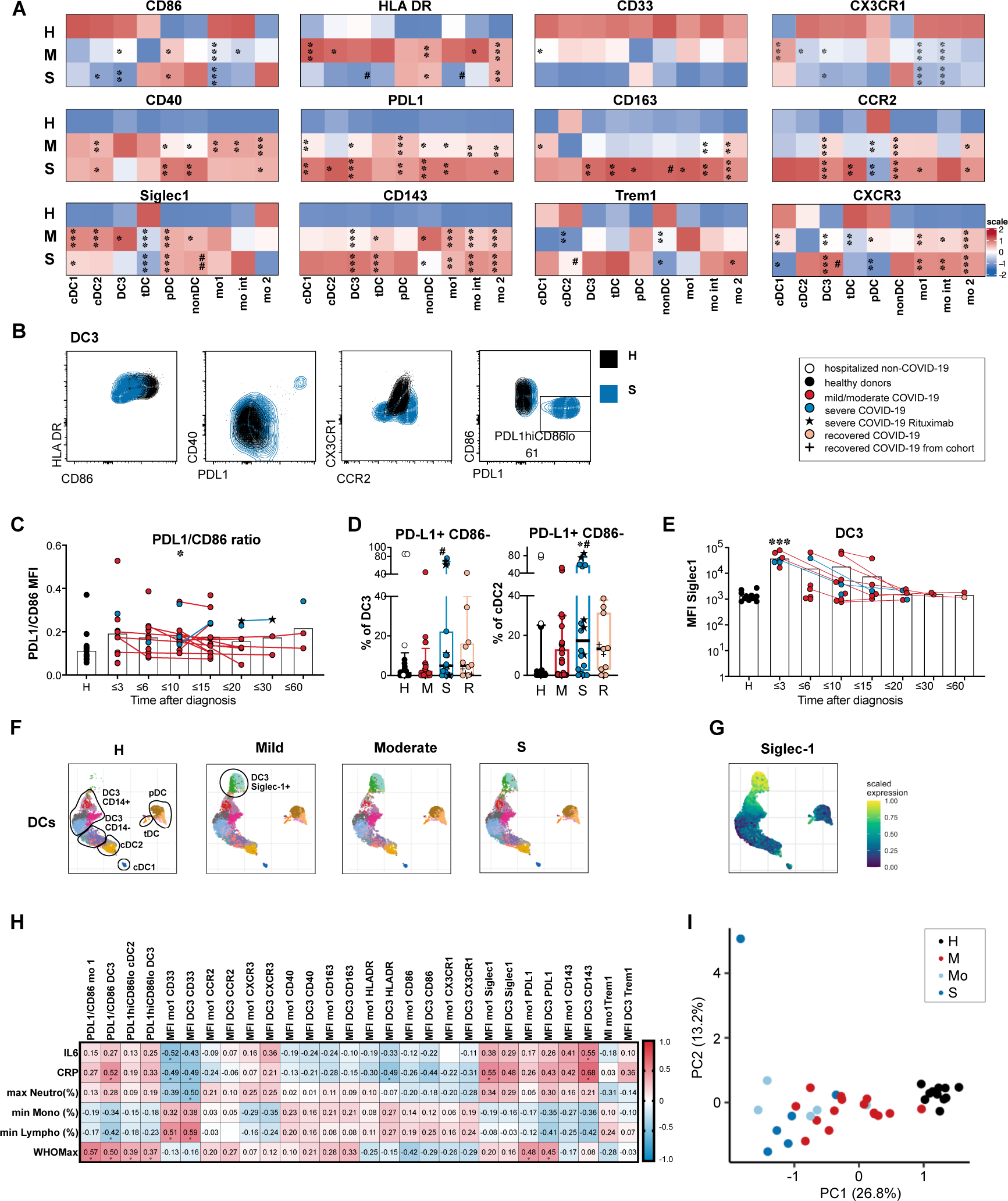
Phenotype alterations in circulating DC and monocyte subpopulations in COVID-19 patients compared to healthy controls. (A) Expression heatmap of log10 transformed median MFI values of surface markers in all DC and monocyte subpopulations in COVID-19 patients with mild/moderate (M, n=20) or severe disease (S, n=6) at the first analysis timepoint compared to healthy donors (H, n=11). The color indicates the scaled expression (z-score standardized) for each cell population (red = high expression, blue = low expression), * significant differences between patients and healthy donors, # significant differences between patients with mild/moderate and severe COVID-19 (ANOVA or Kruskal-Wallis test with Tukey’s or Dunn’s correction for multiple comparisons between H, M and S, p<0.05). (B) Representative results of the expression of HLADR, CD86, CD40, PD-L1, CX3CR1 and CCR2 in DC3 in a healthy control (black) and a patient with severe COVID-19 (blue). (C) Ratio of PD-L1 and CD86 MFI values in DC3 at different grouped time points after diagnosis. Connected lines represent multiple measurements of the same donor at different time points. Columns indicate the mean (n=81, cohorts 2 and 3 combined). (D) Frequency of the PD-L1^hi^ CD86^lo^ population in DC3 and cDC2 in healthy donors (H, black, n=28), non-COVID patients (H, white, n=4) and COVID-19 patients with mild/moderate (M, red, n=39) and severe (S, blue, n=18) disease. (E) Siglec-1 expression (MFI) at different time points after diagnosis in DC3. (F) Clustering analysis was performed on pooled samples of 26 patients and 11 controls. UMAPs of reclustered DCs with Phenograph clusters indicated by colors are shown separately for the indicated groups (same number of cells). DC subpopulations are annotated according to marker expression in phenograph clusters (shown in Fig. S3). (G) Siglec-1 scaled expression indicated by color overlayed on the UMAP embedding (moderate group). (H) Spearman rank correlation coefficients (-1 to 1) for activation markers in mo 1 and DC3 with markers of inflammation and disease severity are shown and indicated by color scale (n=26-41). * adjusted p values below 0.05 (Benjamini-Hochberg procedure) are indicated by asterisks. (I) Principal component analysis using extracted parameters from flow cytometric analysis of all DC and monocyte subpopulations with clinical groups indicated by colors.

Higher expression of the CD163 was detected in monocytes and DC3 of COVID-19 patients with severe disease. TREM-1 was most highly expressed in mo 1 and mo int and significantly upregulated in mo 2 of COVID-19 patients. Expression of CD143 (angioconverting enzyme, ACE) was increased in monocyte subpopulations, cDC2, DC3, and tDCs in COVID-19 patients compared to healthy controls (Fig. 4a), especially at early time points (Fig. S4). ACE2, the primary entry receptor for SARS-CoV-2 was barely detectable on the surface of peripheral blood monocytes and DCs and not induced in COVID-19 patients compared to controls (Fig. S2). CD33 was found to be downregulated in all APC populations of the patients especially in severe disease (Fig. 4a). This may be due to older age, as CD33 was also reduced in older compared to younger healthy donors (Fig. S2). CD143 expression was also influenced by age, but the difference between COVID-19 and healthy controls was more significant than the difference between the age groups (Fig. S2). We did not detect significant differences in expression levels between young and old healthy donors in other markers (Fig. S2).

CCR2 was found to be expressed at higher levels in COVID-19 patients than controls in all monocyte and DC subpopulations except pDC (Fig. 4a and b). The CCR2-CCL2 axis is crucial for the recruitment of inflammatory monocytes to the site of inflammation or infection. It could similarly be involved in the recruitment of DC3, which expressed comparably high levels of CCR2 as classical and intermediary monocytes. CXCR3, which mediates chemotaxis in response to IFN-induced inflammatory chemokines (CXCL9, CXCL10, CXCL11), was also upregulated in COVID-19 patients’ DC3, cDC2 and monocyte subsets, but downregulated in cDC1, tDC, and pDC. CXCR3 expression in DC3 was significantly higher in patients with severe than mild/moderate disease. CX3CR1, which is linked with patrolling ability and survival of monocytes, was downregulated in cDC2, DC3 and monocytes in our patient cohort (Fig. 4a, b). These results show that circulating cDC and monocyte subpopulations in COVID-19 patients are poised to migrate in response to inflammatory chemokine ligands.

Type I IFN-induced Siglec-1 (CD169) was strongly upregulated predominantly in DC3 and mo 1 in the majority of patients sampled until 4 days after diagnosis and in a small subgroup of patients sampled until 15 days after diagnosis, indicating an early transient type I IFN response in most of the patients. We observed rapid downregulation of Siglec-1 expression in longitudinally sampled patients (Fig. 4e). Unbiased mapping of the pooled high-dimensional dataset showed that DC3 were continuously distributed between cDC2 and CD14^+^ monocytes and contributed to a cluster of Siglec-1^hi^ cells, which also contained mo 1 and some mo int (Fig. S3). Reclustering of DCs confirmed the appearance of a separate cluster of CD14^+^ CD163^+^ Siglec-1^+^ DC3 (cl. (cl. 11, 14, 12) in a subgroup of COVID-19 patients (Fig. 4f, 4g and S3). Similarly, a Siglec-1^+^ mo 1 cluster was observed in the monocyte compartment (Fig. S3) showing a similarity of the DC3 and monocyte responses. The Siglec-1^+^ DC3 and mo 1 populations appeared only in samples from COVID-19 patients taken until 14 days after diagnosis and accounted for more than 50 % of the monocytes and DCs in 89% of the samples taken within the first 3 days after diagnosis (Fig. S3). Our deep phenotyping analysis suggests that early transient upregulation of IFN-inducible Siglec-1 occurred irrespective of disease severity. In contrast, the dysregulated PD-L1^hi^ CD86^lo^ HLADR^lo^ activation phenotype of DC3, cDC2 and mo 1 was persisting and more pronounced in severe disease.

### Long-lasting increased proliferative response indicates delayed regeneration of DC and monocyte subsets in COVID-19 patients

Increased myelopoiesis has been described in COVID-19 patients (28, 35).To understand if the altered phenotype of DCs and monocytes was caused by an enhanced recruitment of immature recently generated cells from the bone marrow, we analyzed Ki67 expression as a marker of ongoing or recent 10 proliferation. Even though DCs were reduced in frequency, we found a sizable population of Ki67^+^ cells in all cDC subtypes which tended to be highest in the mild/moderate group (Fig. 5a, b). tDCs and the HLADR^+^ non-DC population had the highest frequencies of Ki67^+^ cells even in healthy/non-CoV controls which was further increased in COVID-19 patients consistent with their precursor/progenitor function (Fig. 5a). The percentage of Ki67^+^ mo 1 was significantly increased in patients with active disease compared to controls (Fig. 5c, d). Increased Ki67 expression was detected in DCs and monocytes of recovered patients and even later than 60 days after diagnosis in some patients indicating enhanced cellular turnover until late timepoints (Figure 5 a-d). The plasma concentrations of FLt3L and GM-CSF, growth factors, which can promote the generation and expansion of DCs and monocytes, were slightly higher in patients compared to heathly controls, especially in those with mild or moderate disease severity (Fig. S4).

**Fig. 5.**
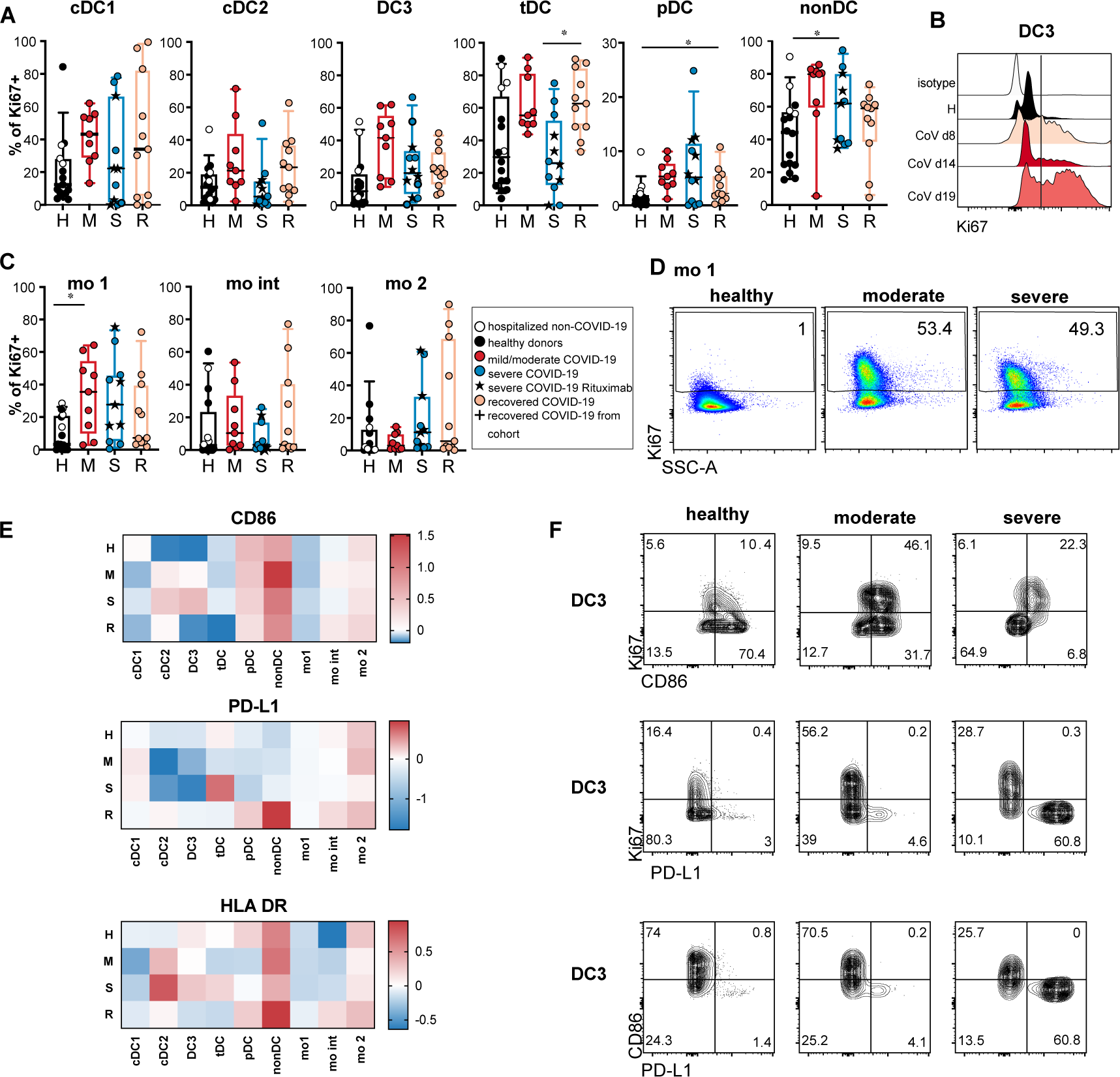
Ki67 expression indicates enhanced and persistent regeneration of DCs and monocytes. (A) Ki67 expression in DC and monocyte subsets was analyzed by intracellular staining and flow cytometry in a subgroup of patients and controls. Healthy donors (H, black, n=12), hospitalized COVID-19 negative patients (white symbols, n=4), acute COVID-19 patients with mild/moderate (M, red, n=9), severe (S, blue, n=12) disease and recovered patients (n=11). (A, C) Percentages of Ki67^+^ cells in each subset are shown (Kruskal-Wallis test with Dunn’s correction, *p<0.05). (B) Representative histogram of Ki67 signal in DC3 in one patient with moderate COVID-19 at different time points after diagnosis. (D) Representative results of Ki67 expression in mo 1 in one healthy, one moderate and one severe patient shown as dot plots. (E) Log2 fold changes of median MFI values of CD86, PD-L1 and HLADR in Ki67^+^ versus Ki67^-^ cells within the indicated populations are shown in the heatmaps indicated by the color scale. (F) Representative results of Ki67, CD86 and PD-L1 expression in DC3 of a healthy control, and 2 COVID-19 patients with moderate and severe disease are shown.

We hypothesized that the unusual phenotype of cDCs with downregulated CD86 (and HLADR in severe cases), and upregulated PD-L1 is caused by enhanced recruitment of immature DCs from BM to blood and should hence be found in the Ki67^+^ fraction. Therefore, we compared the expression of these markers on the surface of Ki67^+^ and Ki67^-^ cells. Remarkably, we found higher expression of CD86 and HLADR and lower expression of PD-L1 in the Ki67^+^ fractions of cDC2 and DC3 (Fig. 5f, g), suggesting that this phenotype alteration in DCs of COVID-19 patients was not caused by the recruitment of immature progenitors from the bone marrow, but could have been induced by external factors such as inflammatory mediators in the blood. We detected increased levels of several cytokines in the patients’ plasma (including IFN-*α*, CXCL10, IFN-*γ*, IL-6, IL-8, CCL2, IL-10, IL-18, IL-23, IL-33) some of which correlated with time after diagnosis (IL-8, IL-23, IL-33) indicating longer-lasting responses (Fig. S4). We found correlations between the plasma concentrations of several of these cytokines and the PD-L1^hi^ CD86^lo^ phenotype of cDC2 and DC3. The strongest correlations (r > 0.4) were found for IFN-*γ*, IL-8, IL-23 and IL-33 (Fig. S4). Thus, prolonged systemic cytokine responses may contribute to the long-lasting phenotypic and functional changes observed in circulating cDCs of COVID-19 patients.

### DC3 and monocytes isolated from the blood of COVID-19 patients show reduced capacity to stimulate naïve CD4^+^ T cells

DC3 have been shown to stimulate naïve CD4^+^ T cells to proliferate and produce IFN-*γ* and IL-17 (33). Due to the observed downregulation of CD86 and upregulation of PD-L1 DC3 and classical monocytes isolated from the blood of COVID-19 patients may be impaired in their ability to stimulate naïve CD4^+^ T cells. DC3 or classical monocytes isolated from COVID-19 patients and healthy controls were cocultured with CFSE-labeled autologous naïve CD4^+^ T cells in the presence of suboptimal TCR stimulation by anti-CD3 antibody. DC3 from COVID-19 patients, which had lower CD86 expression (see Fig. 6e), induced significantly less proliferation and CD69 expression in T cells than DC3 from healthy donors irrespective of glucocorticoid therapy (Fig. 6a, b). Reduced T cell proliferation was also observed in cocultures with monocytes from COVID-19 patients (Fig. 6c). Proliferation and CD69 expression of CD4^+^ T cells in response to stimulation with anti-CD3/CD28 were comparable between patients and controls. Therefore, the reduced T cell response in cocultures with DC3 or monocytes was not due to impaired responsiveness of the patients’ T cells but to the reduced costimulatory activity of DC3 and monocytes. We also detected lower concentrations of the cytokines IL-2, IL-4, IL-5, IL-9, IL-10, IL-13, IL-17A, IFN-*γ*, and TNF-*α* in cocultures of CD4^+^ T cells and DC3 from patients than from controls consistent with the reduced T cell activation. In response to CD3/CD28 stimulation, CD4^+^ T cells from patients produced similar amounts of most of these cytokines and even higher amounts of IL-5 and IL-10, indicating that their ability to differentiate into cytokine-producing Th cells was not generally impaired (Fig. 6d). These results show that phenotypic changes are accompanied by functional impairment of circulating DC3 and monocytes in COVID-19 patients.

**Fig. 6.**
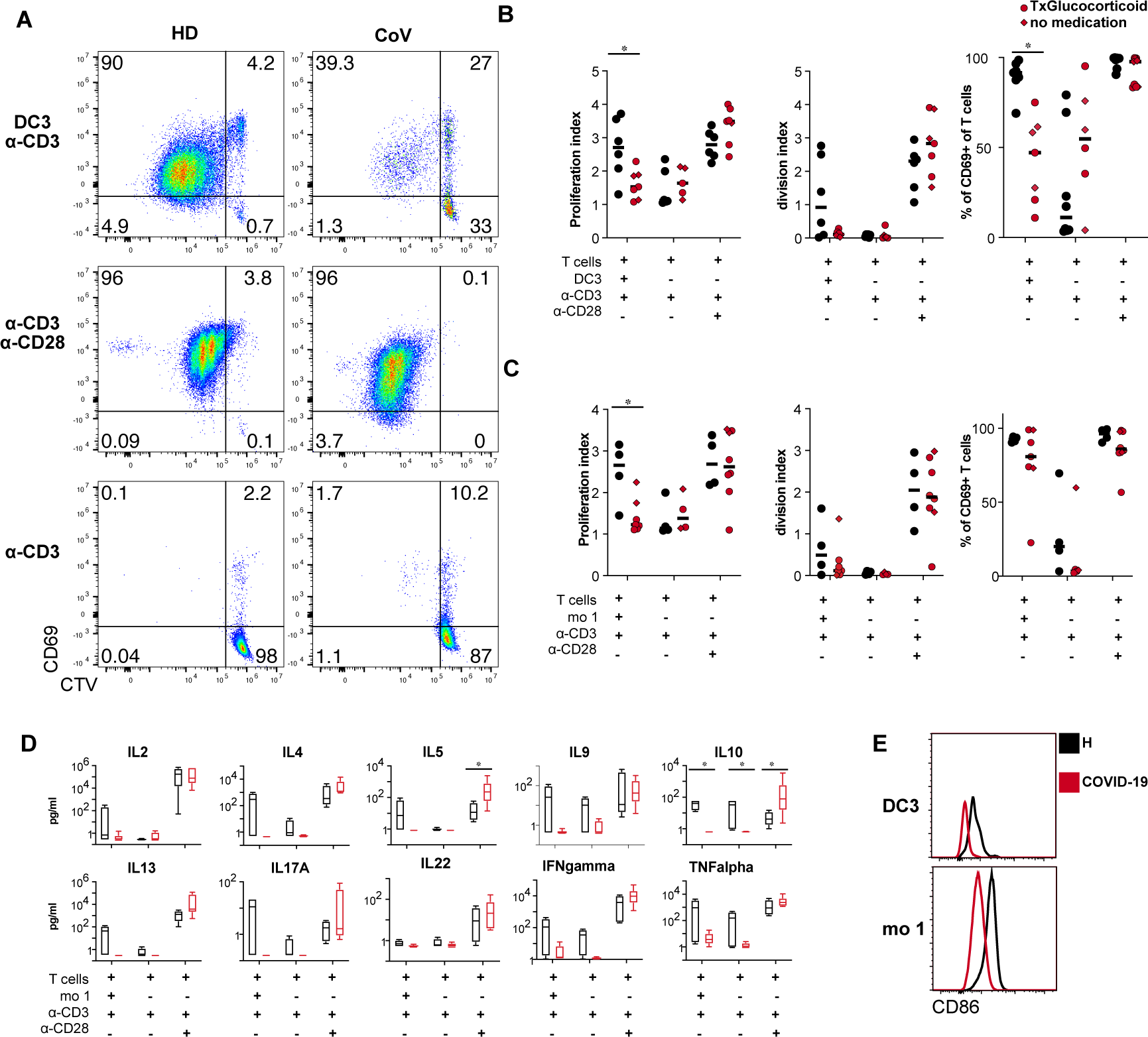
DC3 and monocytes from COVID-19 patients have reduced ability to stimulate naïve CD4 T cells. (A-D) CTV-labeled autologous naïve CD4^+^ T cells were stimulated with immobilized anti-CD3 antibodies in the presence or absence of DC3 and mo 1 sorted from PBMC of healthy donors and COVID-19 patients at a 1:2 ratio for 5 days. Stimulation with anti-CD3/CD28-coated beads was used as a positive control. (A) Representative dot plots showing proliferation of CD4^+^ T cells by CTV dilution and activation by CD69 expression (HD healthy donor, CoV COVID-19 patient). (B) Proliferation index, division index and percentage of CD69^+^ T cells are shown for cocultures with DC3 from healthy controls (black symbols n=6) and COVID-19 patients (red symbols, n=7, circles: glucocorticoid therapy, diamonds: no glucocorticoid therapy). (C) Proliferation index, division index and percentage of CD69^+^ T cells are shown for cocultures with monocytes (H n=4, CoV n=4-7). (D) Cytokines were measured in the supernatants of the experiments with DC3 coculture (H, n=4, CoV n=6). (B, D) *p<0.05, Mann-Whitney test. (E) Representative histogram overlay showing CD86 expression in DC3 and mo 1 sorted from a COVID-19 patient and a healthy donor used for coculture.

### The adaptive immune response is marked by T cell activation and an increase of antibody-secreting cells

Reduced numbers, phenotypic alterations and impaired costimulatory function of circulating DC and monocyte subpopulations found in our patient cohort could affect adaptive immune responses. We, therefore, investigated the frequency of blood T and B cell subpopulations and their activation status. Lymphocyte counts and percentages correlated inversely with disease severity in our patient cohort as expected (see Fig. 1) and the percentages of CD3^+^ T cells were reduced, especially in the group of patients with severe disease (Fig. S5) consistent with T cell lymphopenia. We observed a shift from naïve (CD45RA^+^) to non-naïve (CD45RA^-^) CD4^+^ T cells in the severe group, while the frequency of CXCR5^-^ Th and CXCR5^+^ PD-1^+^ Tfh-like cells within CD4+ T cells was not considerably altered in COVID-19 patients (Fig. 7a and S5). As specific T cell activation in response to acute viral infection can be detected by increased HLADR and CD38 expression (36) we investigated the coexpression of these molecules. The percentage of activated Th and Tfh-like cells was higher in patients with active disease compared to controls and recovered patients (Fig. 7b and c). Increased activation was observed in samples taken until 30 days after diagnosis (Fig. 7d). The CXCR3^-^ CCR6^-^ Th0/2 cell fraction was increased in patients with severe disease but contained only a low percentage of activated cells. Circulating Th and Tfh-like cells expressing CXCR3 and or CCR6 showed increased activation in patients with active COVID-19 (Fig. S5). In the CD8^+^ T cell compartment, we observed a reduction of CD45RA^+^ CD27^+^ naïve CD8^+^ T cells with a concomitant increase in CD45RA^-^ CD27^+^ CD8^+^ T cells containing T_CM_ (Fig. 7e and S5). CD8^+^ T cell activation, which was detected mainly in the T_CM_ and T_EM_ containing fractions was increased in patients with active COVID-19 *vs* controls and recovered patients (Fig. 7e-h). The highest frequencies of activated CD8^+^ T cells were observed between 6 and 15 days after diagnosis (Fig. 7g).

**Fig. 7.**
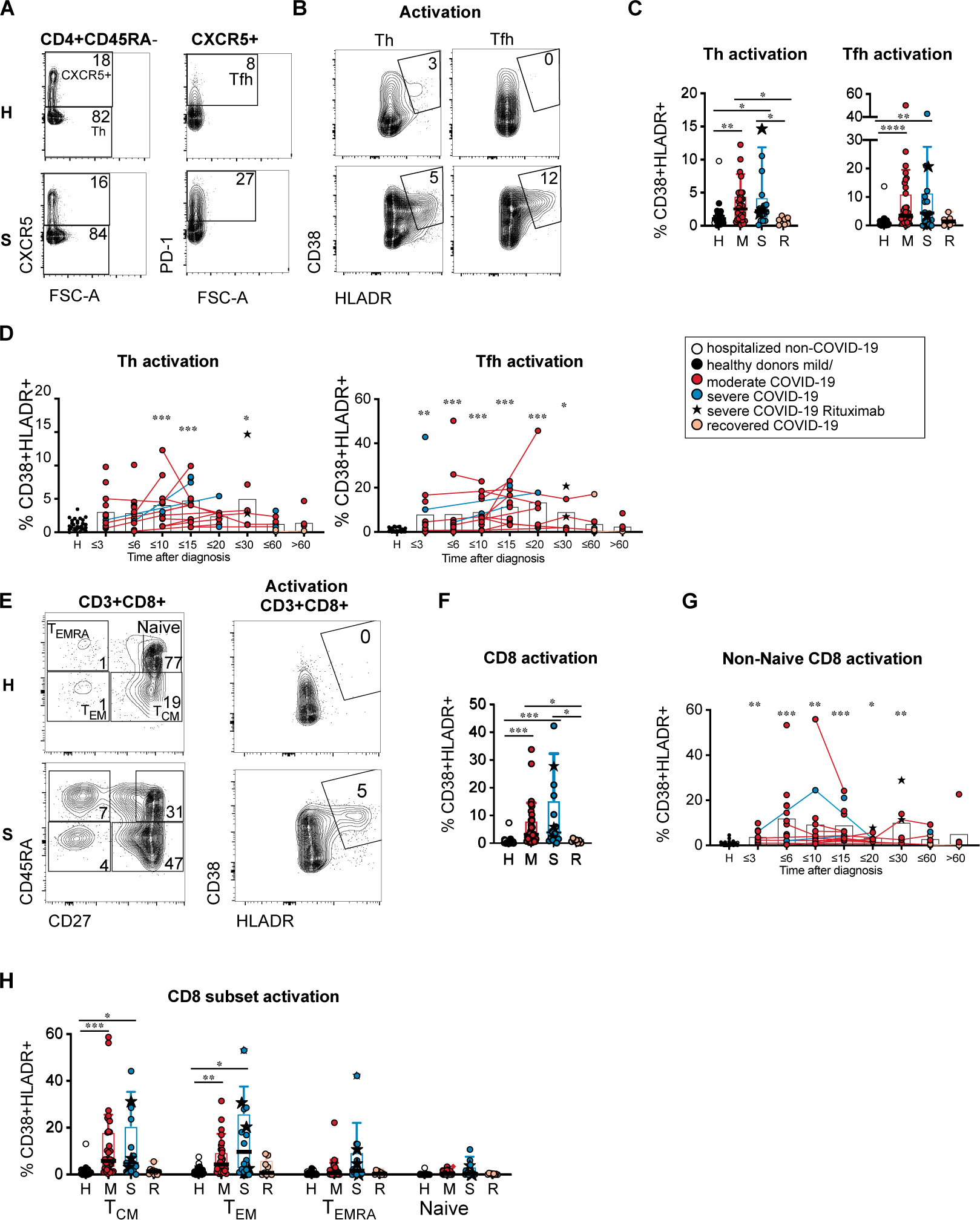
Heterogeneous T cell activation in COVID-19 patients. (A and B) Representative results of one healthy donor and one COVID-19 patient with severe disease are shown. Numbers indicate percentages. (A) CXCR5 and PD-1 expression in CD3^+^ CD4^+^ CD45RA^-^ CD25^lo/int^ T cells and gating of Tfh-like cells. (B) CD38 and HLADR expression in CXCR5^-^ Th and CXCR5^+^ PD-1^+^ Tfh-like cells. (C) Percentage of activated cells within Th and Tfh-like cells in healthy/non-COVID controls (H, n=24) and acute COVID-19 patients with mild/moderate (M, n=35) or severe disease (S, n=16) at the first analysis timepoint and recovered patients (R, n=7). (D) Percentages of activated T cells within the Th and Tfh-like cells at different time points (days) after diagnosis of SARS-CoV-2 infection. Connected lines represent multiple measurements of the same donor at different time points (n=103). Columns indicate the mean. (E) Representative results of one healthy donor and one COVID-19 patient with severe disease. Left: CD8^+^ naïve and memory subsets according to CD27 and CD45RA expression. Right: CD38 and HLADR expression in CD8^+^ T cells. Numbers indicate percentages. (F) Percentage of activated cells within CD8^+^ T cells in the indicated groups (as in C). (G) Percentage of activated cells within non-naïve CD8^+^ T cells at different time points after diagnosis (n = 105, shown as in D). (H) Percentages of activated T cells within naïve and memory CD8^+^ T cell subsets in the indicated groups (as in C). (C, D, F, G, H) Kruskal-Wallis test with Dunn’s correction, *p<0.05, **p>0.01, ***p<0.001. (D, G) comparison of grouped timepoints to the control group.

B cell frequencies were similar in patients and controls, but the percentage of CXCR5^+^ B cells was significantly reduced in active COVID-19 (Fig. 8a and b). We detected decreased naïve and memory but increased class-switched memory B cells compared to controls in a subgroup of COVID-19 patients (Fig. 8c and g). Antibody secreting cells (ASC) were expanded in some but not all of the patients and returned to the level of healthy controls in the recovered patients (Fig. 8d and g). The expansion of ASC was already seen at early time points and persisted until 20 days after diagnosis and even longer in some patients (Fig. 8e). Anti-SARS-CoV-2 spike S1 IgG antibodies were detected in 70.9% and anti-SARS-CoV-2 nucleocapsid IgG antibodies in 77.8% of patients at the latest available timepoint (n=54-55) and in 90.3 % and 93.5 % of patients sampled later than 15 days after diagnosis (n=32) indicating specific antibody production in the majority of patients (data not shown). Antibody levels increased with time after diagnosis and in longitudinally sampled patients (Fig. 8f). The frequency of activated Tfh-like cells correlated only weakly and the frequency of ASC did not correlate with anti-SARS-CoV-2 antibody levels (Fig. S5).

**Fig. 8.**
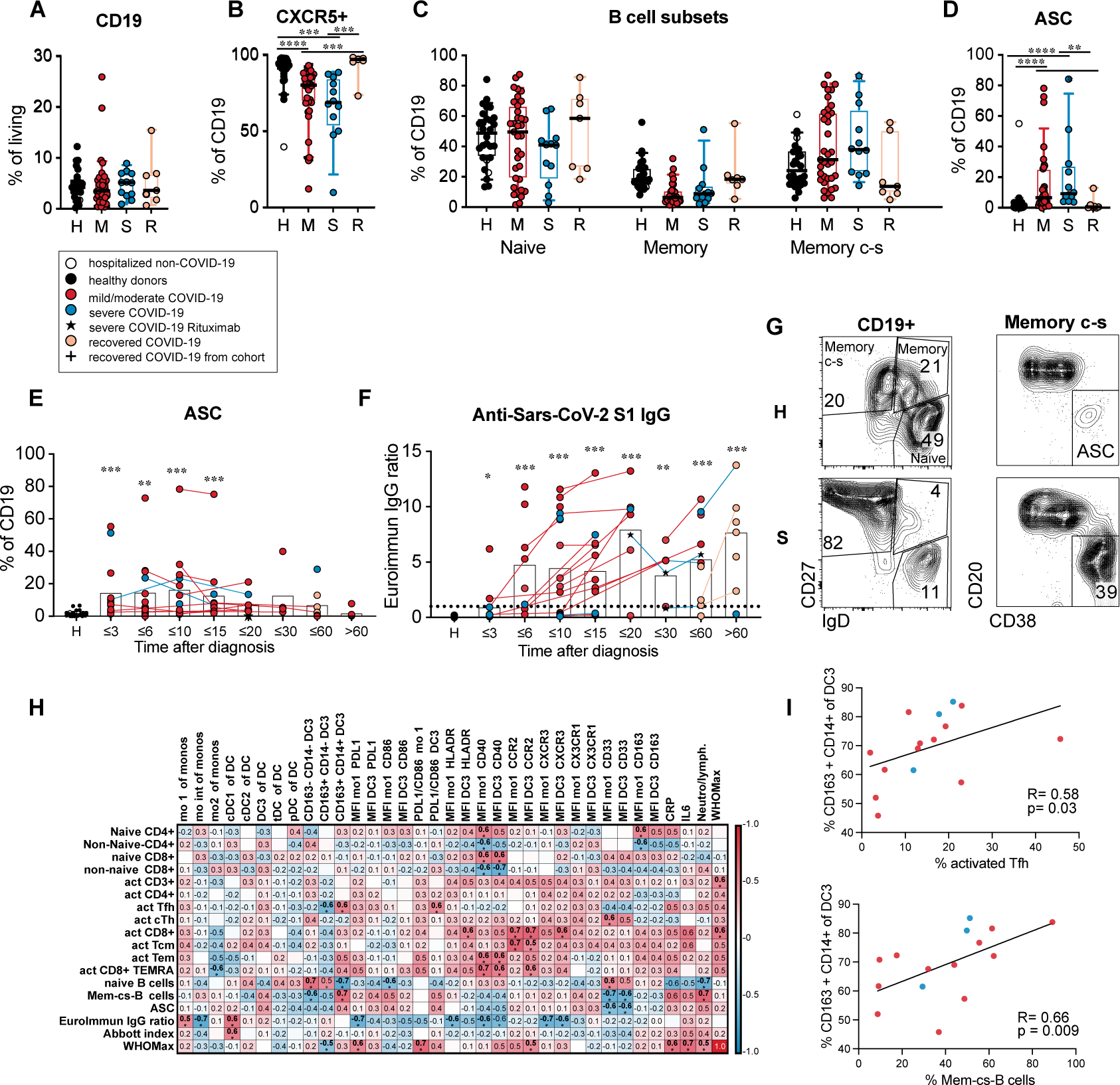
DC3 and monocyte phenotype correlates with Tfh and B cell activation. (A) Frequency of CD19^+^ B cells within living PBMC, (B) percentage of CXCR5^+^ cells within the CD19^+^ B cells, (C) percentage of naïve and memory B cell subsets within CD19^+^ cells, (D) percentage of antibody-secreting cells (ASC) within CD19^+^ B cells in healthy/non-COVID controls (H, black/white symbols n=24) and acute COVID-19 patients with mild/moderate (M, red, n=35) or severe disease (S, blue, n=13) at the first analysis timepoint and in recovered patients (R, orange, n=7). Patients that had received B cell-depleting therapies were excluded. Kruskal-Wallis test with Dunn’s correction. (E) Frequency of ASC within CD19^+^ B cells (n=102), (F) anti-SARS-CoV-2 spike S1 IgG levels (Euroimmun IgG ratio) in plasma samples (n=105) of patients at different time points and in controls. The dotted line indicates the cutoff value for the antibody test. (E, F) Symbols indicate individual measurements (see legend in A). Connected lines represent multiple measurements of the same donor at different time points. Columns indicate the mean. Data from each grouped time point were compared with data from the control group. Kruskal-Wallis test with Dunn’s correction, *p<0.05, **p>0.01, ***p<0.001. (G) Gating strategy for B cell subpopulations and ASC. Representative results from a healthy donor and a COVID-19 patient with severe disease are shown. (H) Correlation analysis of innate parameters up to 10 days after diagnosis (horizontal) with adaptive parameters at day 10 to 25 after diagnosis (vertical) in the same patients (n=9-17). Spearman’s rank correlation coefficients (-1 to 1) are indicated by the color scale. * adjusted p values below 0.05. (I) Correlation analysis of the percentage of CD163^+^ CD14^+^ cells within DC3 (until day 10) and the percentage of activated cells within Tfh-like cells or the percentage of class-switched memory B cells within CD19^+^ cells (day 10-25 after diagnosis, n=15). Spearman’s rank correlation coefficient, p-value and linear regression line are shown.

To better understand the connection between the innate and the adaptive immune response, we performed a correlation analysis of innate parameters in the early phase (day 0-10) and adaptive parameters in the later phase (day 10-25 after diagnosis) in longitudinally sampled patients (Fig. 8h). The expression of CCR2, CXCR3, HLADR and CD40 in DC3 and monocytes correlated with subsequent CD8^+^ T cell activation and inversely with anti-S1 antibody levels indicating that this APC phenotype could be relevant for CD8^+^ T cell activation in response to SARS-CoV-2 infection. The frequency of CD163^+^ CD14^+^ cells within DC3 and the PD-L1/CD86 ratio in DC3 correlated positively with the frequency of activated Tfh cells and cs-mem B cells (Fig. 8i) and ASC, but not with antibody titers (Fig. 8h). This DC3 phenotype, as well as Tfh and B cell activation, also correlated with increased inflammatory markers (CRP, IL-6), neutrophil/lymphocyte ratio and disease activity (Fig. 8h, see also Fig 3f and 4h). Thus, the PD-L1^hi^ CD86^lo^ CD163^+^ CD14^+^ differentiated DC3 phenotype and subsequent Tfh and B cell activation are linked to the systemic inflammatory response and lymphopenia characteristic of more severe disease.

## Discussion

In this study, we provide an in depth characterization of DC and monocytes subpopulations in the blood of hospitalized patients with mild, moderate or severe COVID-19 compared with healthy controls. Changes in DC/monocyte composition and phenotype were connected with parameters of inflammation and activation of adaptive immunity. We found that all DC subpopulations were profoundly and persistently depleted from the blood in COVID-19 patients while Lineage^-^ HLADR^+^ cells lacking DC markers expanded. Correlating with systemic inflammation, DC3 contained more CD163^+^ CD14^+^ cells. Similar to classical monocytes, DC3 showed dysregulated activation with low CD86, high PD-L1 and CD40 expression and impaired ability to stimulate T cells. The long-lasting proliferative response indicated increased turnover and delayed regeneration of the DC and monocyte compartment. Thus, reduced APC numbers and functionality may contribute to an immunosuppressed state in COVID-19 patients, making them vulnerable to other infections or virus reactivation.

The long-lasting reduction of all DC subsets in the blood which occurred in patients with mild/moderate and severe disease, was accompanied by increased proliferation, which - although detectable for a long time after diagnosis - did not fully restore the circulating DC compartment. The reduction in DCs, which has also been observed in previous studies (16, 18, 19, 37) may be due to increased emigration from the blood and sequestration in tissues, auch as the inflamed lung or lymphoid tissues. We found that similar to monocytes CCR2 and CXCR3 were upregulated in DC3 of COVID-19 patients suggesting that DC3 together with monocytes may be recruited from the circulation to the infected lung in response to a gradient of CCL2 and CXCR3 ligands CXCL9/10/11 (38). Indeed inflammatory chemokines CCL2, CCL3 and CCL4 have been found at higher concentrations in the airways compared to the plasma in patients with severe COVID-19 (24). cDC2 may follow a similar route, while cDC1 did not upregulate these receptors and pDCs even showed downregulation of CCR2 and CXCR3. This is consistent with preferential recruitment of cDC2 versus cDC1 to the lung (18) and low numbers of pDCs found in the airways and lungs of COVID-19 patients (38, 39). cDC1 and pDC could be reduced due to sequestration in lymphoid tissues or enhanced cell death as shown for pDCs (40). Reduction of circulating DCs due to productive infection by SARS-CoV-2 is unlikely. We did not detect expression of the major entry receptor ACE2 on blood DC and monocytes in accordance with previous reports (41, 42).

While differentiated DC subsets were reduced, we found that Lineage^-^ HLADR^+^ CD86^+/–^ CD45RA^+/–^ proliferating cells lacking typical DC markers were greatly expanded in the blood of COVID-19 patients. Their phenotype did not overlap with that of previously described DC or monocyte/macrophage or lymphoid progenitor populations. Expression of HLADR and lack of CD33, CD14 and CD15 expression indicated that they are not typical myeloid derived suppressor cells. It is unlikely that these cells were activated proliferating innate lymphoid cells or precursors due to lack of CD127 expression. A similar immature HLADR^+^ cell type with poor antigen-presenting capacity and a low response to TLR stimulation was found to be expanded at the expense of cDCs and pDC in the blood of patients with cancer or acute malaria (43, 44). The long duration of cDC reduction and immature HLADR^+^ cell expansion in the blood of COVID-19, even in convalescent patients, indicated delayed regeneration of the DC compartment. The “non-DCs” described in our study could be an immature DC-like population appearing in COVID-19 due to hyperinflammation and increased myelopoiesis.

We observed a shift towards a more mature CD163^+^ CD14^+^ phenotype within the DC3 subset in acute COVID-19 correlating with disease severity and inflammatory markers. A similar change in DC3 phenotype has been observed in the blood of SLE patients and in melanoma patients coinciding with inflammation (33, 45). CD163^+^ CD14^+^ DC3 have been shown to express higher levels of proinflammatory genes, secrete more proinflammatory mediators and induce Th17 polarization more efficiently than CD163^-^ CD14^-^ DC3 (33). It is still unclear, however, if the CD163^+^ CD14^+^ phenotype of DC3 in the blood of COVID-19 patients contributes to or is a byproduct of the inflammatory response.

Clusters of monocytes and DC3 with high expression of Siglec-1 appeared in the blood of COVID-19 patients at early timepoints and disappeared at later time points, indicating a robust but transient type I IFN response. Consistent with this dynamic expression pattern Siglec-1 was shown to serve as a negative feedback regulator of type I IFN production in response to viral infection (46). Siglec-1^+^ expression on monocytes was a promising biomarker for the early 19 diagnosis of COVID-19 in the emergency room (47). In contrast to this study, we detected high Siglec-1 expression in monocytes and DC3 only in half of the patients analyzed, most likely due to later sampling timepoints. We also found ACE/CD143 to be upregulated at early timepoints in monocytes and DC3 of COVID-19 patients correlating with markers of inflammation. Increased expression of this carboxypeptidase has been observed in bacterial infections but not typically in viral infections (48). ACE/CD143 was found to promote TNF-*α* and IL-6 production, adhesion and transmigration of myeloid cells in response to CCL2 (49) and could therefore be involved in tissue migration and cytokine response of monocytes and DC3 in COVID-19 patients.

The expression of costimulatory molecules was differentially regulated in different blood APC subsets. In pDCs, we observed increased expression of CD86, CD40 and PD-L1, but did not detect diversification into distinct activated pDC effector subsets as described by Onodi et al. after exposure to SARS-CoV-2 *in vitro* (41). DC3, cDC2 and mo 1 showed reduced CD86 expression and increased CD40 and PD-L1 expression in the patients, most pronounced in severe disease. Reduced expression of CD86 in circulating monocytes and cDCs has been described as a feature of severe COVID-19 (16, 19, 20) but was also found in patients with less severe disease in our study. We observed reduced HLADR expression on monocytes and cDCs only in patients with severe COVID-19 and increased HLADR levels in monocytes in a subgroup of patients with mild disease consistent with published data (28). The frequency of proliferating DCs and monocytes was increased in the patients of our cohort in line with increased myelopoiesis (28, 35), but only Ki67^-^ DCs showed the 20 PD-L1^hi^ CD86^lo^ HLADR^lo^ phenotype and proliferating DCs had a similar phenotype as healthy donors. Therefore, it is unlikely that the phenotypic and functional alterations were due to impaired differentiation of DCs from precursors. Instead, the observed changes may be caused by circulating inflammatory mediators. Correlation of the PD-L1^hi^ CD86^lo^ HLADR^lo^ phenotype in monocytes and cDCs with the plasma levels of CRP, IL-6 and other proinflammatory cytokines supported this assumption.

The dysregulated phenotype of DC3 and classical monocytes translated into a defect in their ability to support efficient proliferation and differentiation of naïve CD4^+^ T cells. The reduced T cell proliferation observed in coculture with APCs from COVID-19 patients was not due to impaired responsiveness of the T cells. It was shown that DCs isolated from the blood of patients with severe COVID-19 are less responsive to stimulation with TLR ligands and cytokines, further supporting their impaired functionality (19, 20). Monocytes isolated from the blood of COVID-19 patients were even shown to actively suppress T cell activation (50).

The inability of DCs to sufficiently prime T cell responses could have dire consequences in COVID-19 patients leading to inadequate adaptive immune responses against SARS-CoV-2, delaying clearance of the virus. However, we found T cell activation and SARS-CoV-2-specific antibody production in most patients in our cohort. The frequency of activated T cells in our cohort was highly variable and a subgroup of patients lacked T cell activation above that of healthy controls. This observation is consistent with findings by others (17, 27, 21 51, 52) and was also seen for SARS-CoV-2 specific T cells (12, 53), but contrasts with responses seen in other acute viral infections or vaccinations (36, 54). It may reflect insufficient T cell activation or preferential activation of T cells recruited to the airways compared to circulating T cells (24).

The CD14^+^ CD163^+^ PD-L1^hi^ CD86^lo^ phenotype of DC3 correlated with the activation of circulating Tfh cells, the frequency of class-switched B cells and ASC and with markers of inflammation, but not with anti-SARS-CoV-2 antibody levels. It remains to be investigated if dysregulated activation of DCs directly influences activation of Tfh cells and B cells or if both are affected by the prolonged systemic inflammatory response in COVID-19 patients. Altered DC phenotype and function may contribute to the observed lack of coordination between T cell activation and antibody responses in COVID-19 patients that was similarly shown for SARS-CoV-2-specific T cells and neutralizing antibodies (12).

In summary, we provide evidence that the depletion of circulating DCs, delayed regeneration and phenotypic alteration are long-lasting effects of COVID-19 infection. The persistent phenotypic alteration and dysfunctionality of circulating DCs and monocytes was especially apparent in more severe disease and associated with the prolonged inflammatory response. The consequences of depletion and dysfunctionality of blood APCs are not known. While these changes may reflect a regulatory mechanism to reduce overactivation of the immune response in COVID-19, the described long-lasting alterations together with the profound lymphopenia could make patients more vulnerable to secondary infections, which were shown to be more prevalent in COVID-19 patients (55, 56). This needs to be taken into account in the clinical management of COVID-19.

## Materials and methods

### Patients and healthy controls

Patients are part of the COVID-19 Registry of the LMU University Hospital Munich (CORKUM, WHO trial id DRKS00021225). In the framework of the CORKUM biobank, blood samples were collected from *≥* 18 yrs patients who were diagnosed with COVID-19 by positive SARS-CoV-2 PCR result between March 2020 and January 2021 at LMU Klinikum and had consented to biobanking. PBMC, plasma, and serum were prepared and cryopreserved.

From this biobank, cryopreserved PBMC samples of 26 patients were selected, of whom the first sample had been taken within 3 weeks after the date of the positive PCR result (cohort 2). Of the 26 patients, 23 patients were hospitalized and 3 patients were diagnosed in the ER and discharged home. From 13 patients, only one timepoint could be obtained, which was taken between 0 and 17 days after diagnosis. From 13 patients, longitudinal samples from 1-3 additional time points were analyzed. As a control for this cohort, we used cryopreserved PBMC of 11 healthy donors aged 22 – 54 yrs prepared from leucocyte reduction chambers after thrombocyte donations. To check for age effects, samples from another cohort of patients (cohort 3) were thawed and analyzed, consisting of 15 COVID-19 patients aged between 38-87 years and 8 age-matched healthy controls aged between 56-81 years (H2) as well as younger healthy controls aged between 22-54 (H, same donors as from cohort 2). Additionally, we obtained fresh blood samples from COVID-19 patients diagnosed and treated at LMU Klinikum since mid-May 2020 and used freshly isolated PBMC for flow cytometric analysis (cohort, 4 n=29; and cohort 5, n=19). PBMC freshly prepared from healthy blood donors (hospital and laboratory workers) and from patients who were hospitalized for other reasons and tested negative for SARS-CoV-2 were used as controls. The clinical and laboratory data of each cohort are described in supplementary table 1. All clinical and routine laboratory data were collected and documented by the CORKUM study group. An ordinal scale adopted from the WHO (31) was used to grade disease severity. 1: no limitations of activity; 2: limitations of activity, 3: hospitalized, no oxygen; 4: oxygen by mask or nasal tube; 5: non-invasive ventilation; 6: invasive ventilation; 7: organ support (extracorporeal membrane oxygenation); 8: death. Using the maximal score reached (WHO max) mild (1–3), moderate (4–5) and severe disease (6–8) were distinguished. Patients were classified as recovered when discharged with *≤* WHO score 2 and > 21 days after diagnosis. Immune cell population frequencies from cohorts 2, 3 and 4 were summarized. Summary cohort 1 included 31 COVID-19 negative controls (median age 42, range 22-81), 39 mild/moderate COVID-19 patients (median age 58, range 27-89), 18 severe COVID-19 patients (median age 71, range 40-87) and 11 recovered patients from which 3 were already analyzed during acute disease (median age 56, range 25-88). Five patients had received B cell depleting therapy (Rituximab) within 4 weeks before the diagnosis. These patients all had severe disease manifestations. These patients were excluded for analysis of B cell subpopulations. Detailed clinical characteristics and laboratory parameters for each cohort are shown in table S1.

### Sample preparation

Research blood samples were collected in serum and lithium-heparin tubes and processed within 6 hours after venipuncture. Plasma and serum were separated by centrifugation and cryopreserved at -80C. Peripheral blood mononuclear cells (PBMCs) were isolated by Ficoll density gradient centrifugation and either used directly or resuspended in 90% heat-inactivated FCS/10 % DMSO (v/v) to be stored in liquid nitrogen.

### Flow Cytometry

Cryopreserved PBMC samples of patients and controls were thawed, processed, stained, and analyzed by flow cytometry together. Freshly isolated PBMC from COVID pts and COVID-negative controls were stained and analyzed together on the day of sampling. PBMC stained in 50µl of PBS, 2mM EDTA, 10% FCS (v/v) containing FcR blocking reagent (Miltenyi Biotec) with fluorescently labeled antibodies as indicated in table 2 and incubated for 30 min at 4°C. Fixable viability dyes were used according to the manufacturers’ protocol. Cells were fixed with BD Cytofix (Cat. # 554655), washed and resuspended in PBS. Intracellular staining for Ki67 was performed using the Transcription Factor Staining Buffer Set (ThermoFisher, cat. # 00-5523-00) following the manufacturers instructions. Samples were measured using the Cytek Aurora (Cytec Biosciences) with the recommended Cytek assay settings, where gains are automatically adjusted after each daily QC based on laser and detector performance to an optimal value, ensuring comparability between measurements. Cells from co-culture experiments were measured using the Cytoflex S flow cytometer (Beckman Coulter). FCS files were exported and analyzed with FlowJo software v10.7.1.

### Cell isolation and culture

Cells were sorted from PBMC using the BD FACSAria™ Fusion (BD Biosciences). For the T cell coculture, DC3 (HLADR^+^, CD88/89^-^, CD16^-^, CD56^-^, CD66b^-^, CD15^-^, CD4^-^, CD8^-^, CD11c^+^, CD5^-^, CD1c^+^), classical monocytes (HLADR^+^ CD88/89^+^, CD14^+^, CD16^-^, CD56^-^, CD66b^-^, CD15^-^) and naïve CD4^+^ T cells (CD4^+^, CD45RA^+^, CD8^-^) were sorted. T cells were stained with Cell Trace Violet dye (ThermoFisher, cat. #C34557) washed twice with RPMI 1640 (10% FCS) and cocultured with DC3 or monocytes (APC:T 1:2 ratio) in 150 µl of RPMI 1640 (Biochrom, 10% FCS, 100 U/ml penicillin, 100 μg/ml streptomycin, 1% non-essential amino acids, 1 mM sodium pyruvate, 2 mM GlutaMAX™, 0.05 mM β-mercaptoethanol) on a 96-well flat bottom plate coated with anti-CD3 antibody (10µg/ml, cat. # 317325, BioLegend). 7 x 10^3^ DC3 or 5 x 10^4^ monocytes were used per well. Human T-Activator CD3/CD28 Dynabeads™ (ThermoFisher, cat. #111.61D) were used as a positive control stimulus. After 5 days, cells were harvested and measured using the CytoFLEX S flow cytometer (Beckman Coulter). Supernatants were collected and stored at -20°C.

### Cytokine detection by ELISA and cytometric bead array

Cytokines were measured in plasma samples using the LEGENDplex™ human inflammation assay (cat. # 740809, BioLegend) according to the manufacturer’s instructions. CXCL10/IP-10 was measured by Elisa (cat. # 550926, BD) using 1:20 diluted plasma. FLT3L ELISA (cat. # DY308, R&D) and GMCSF ELISA (cat. # 555126, BD) were performed with 1:2 diluted plasma. Coculture supernatants were measured using the LEGENDplex™ T helper assay (cat. # 741028, BioLegend).

### Clustering analysis of flow cytometric data

Data was processed with R/bioconductor. Unless stated otherwise, default parameters for function calls were used. FlowJo workspace was imported with flowWorkspace::open_flowjo_xml (flowWorkspace version 4.2.0). Cells passing the “HLADR+ Lin-“ gate were selected for further analysis. Cells with negative FI and failing upper boundary filtering on all features except Axl, Siglec and CCR2 were removed. Finally, data was subsampled to 35000 cells per sample (flowCore::filter, version 2.2.0) and converted to a SingleCellExperiment using CATALYST::prepData (version 1.14.0) with parameters FACS=T and cofactor=150 for arcsine transformation.

First clustering was performed on features CCR2, CD163, HLADR, CD16, CD86, CD14, CD141, CD123, Axl, Siglec1, CD88 89, CD5, CD1c with Rphenograph (version 0.99.1). After removal of contaminants, cells were reclustered using the same features, functions and parameters. Data was visualized using CATALYST functions (Crowell H, Zanotelli V, Chevrier S, Robinson M (2020). CATALYST: Cytometry dATa anALYSis Tools. R package version 1.14.0, https://github.com/HelenaLC/CATALYST).

### Anti-SARS-CoV-2 antibody detection assays

The following commercial CE in vitro diagnostics (IVD) marked assays were used to determine the presence of SARS-CoV-2-specific antibodies in serum specimens: Architect SARS-CoV-2 IgG (6R86, Abbott, Illinois, USA) detecting anti-nucleocapsid antibodies and Anti-SARS-CoV-2-ELISA IgG (EI 2606-9601 G, EuroImmun, Lübeck, Germany) recognizing antibodies against the S1 domain of viral spike protein. Assays were performed in accordance with the manufacturers’ instructions by trained laboratory staff on appropriate analyzers and with the specified controls and calibrants, using thresholds for calling positives, indeterminates and negatives set by the manufacturers.

### Statistical analysis

Statistical analysis was performed using GraphPad Prism 9.1.0 and R 4.0.3 (packages used: ggplot2_3.3.3, ComplexHeatmap_2.4.3, ggstatsplot_0.6.8, bestNormalize_1.7.0, robustbase_0.93.7). Box plots show the 25 to 75 percentile, whiskers show the 10 to 90 percentile, horizontal lines indicate the median. Normality was tested using the Shapiro-Wilk test. Normally distributed data was tested with an ANOVA and not normally distributed data with the Kruskal-Wallis test. Multiple testing was corrected using the Tukey’s or Dunn’s multiple comparison test. Subpopulations containing less than 10 cells were excluded from analysis. p-values below 0.05 were considered to indicate statistically significant differences. Spearman correlation coefficients were calculated and the Benjamini Hochberg procedure was used to correct for multiple testing and control the false discovery rate. Samples from patients with B cell depleting therapy were excluded for B cell analysis and correlations.

### Study approval

The study was conducted in the framework of the COVID-19 Registry of the LMU University Hospital Munich. Written informed consent was received from participants prior to inclusion in the study and patient data were anonymized for analysis. The study was approved by the local ethics committee (No. 20-245). Additional approval was obtained for the analyses shown here (No. 592-16) and for the use of blood samples from healthy donors (No. 18-415).

## Author contributions

Conceptualization, A.B.K., E.W., M.S.

Investigation, E.W., L.Ri., L.W., A.R., A.L., L.Ra., J.K., P.R.W.

Formal Analysis, E.W., L.Ri., K.L., A.B.K., P.R.W., T.S., C.S.

Methodology, E.W., L.Ri., A.R., A.L., P.R.W., J.E.H., D.B., S.R.

Resources, J.C.H., C.S., M.M., K.S., M.B-B

Supervision, A.B.K., M.S., T.B., O.T.K, J.C.H., C.S., M.M.

Funding Acquisition, A.B.K., M.S., T.B., O.T.K, M.B-B.

Visualization, E.W., L.Ri., K.L.

Writing - Original Draft, E.W., A.B.K.

Writing – Review & Editing, E.W., A.B.K., T.B., D.B., S.R.

## Acknowledgments

This work is part of the theses of Elena Winheim and Linus Rinke. We would like to thank all CORKUM investigators and staff. The authors thank the patients and their families for their participation in the CORKUM registry. We would like to thank Patricia Späth for assistance in sample preparation and Yvonne Schäfer for technical assistance. We acknowledge the Core Facility Flow Cytometry of the Biomedical Center, LMU Munich and thank Lisa Richter and Pardis Khosravani. A.K. is supported by the Deutsche Forschungsgemeinschaft under SFB1054-TPA06 (210592381), SFB/TR237-B14 (369799452) and KR2199/10-1 (391217598), and received funding from the Bavarian State Ministry of Science and the Arts. E.W. received a scholarship from the Villigst Foundation. T.B. is supported by the Deutsche Forschungsgemeinschaft under SFB1054/TPB03 (210592381). M.S. was supported by the Deutsche Forschungsgemeinschaft under SFB 1243-A10 (278529602) and SU197/3-1 (451580403), the Bavarian Elite Graduate Training Network and the Wilhelm Sander Stiftung (project no. 2018.087.1). A.R. was supported by the Else-Kröner-Fresenius-Stiftung. D.B. was supported by Deutsche Forschungsgemeinschaft under Emmy Noether Programme BA 5132/1-2 (252623821), SFB1054/TPB12 (210592381), and Germany’s Excellence Strategy EXC2151 (390873048). S.R. was supported by the Deutsche Forschungsgemeinschaft under Ro 25257/-1 (391217598) and SFB/TR-237-B14 (369799452). The CORKUM biobank is funded, in part, by the Federal Ministry of Education and Research (BMBF) initiative “NaFoUniMedCovid19” (01KX2021), LMUexcellent, the Free State of Bavaria under the Excellence Strategy of the Federal Government and the States, and the Faculty of Medicine of the LMU München.

