## Supplementary Figures for "Impaired function and delayed regeneration of dendritic cells in COVID-19"

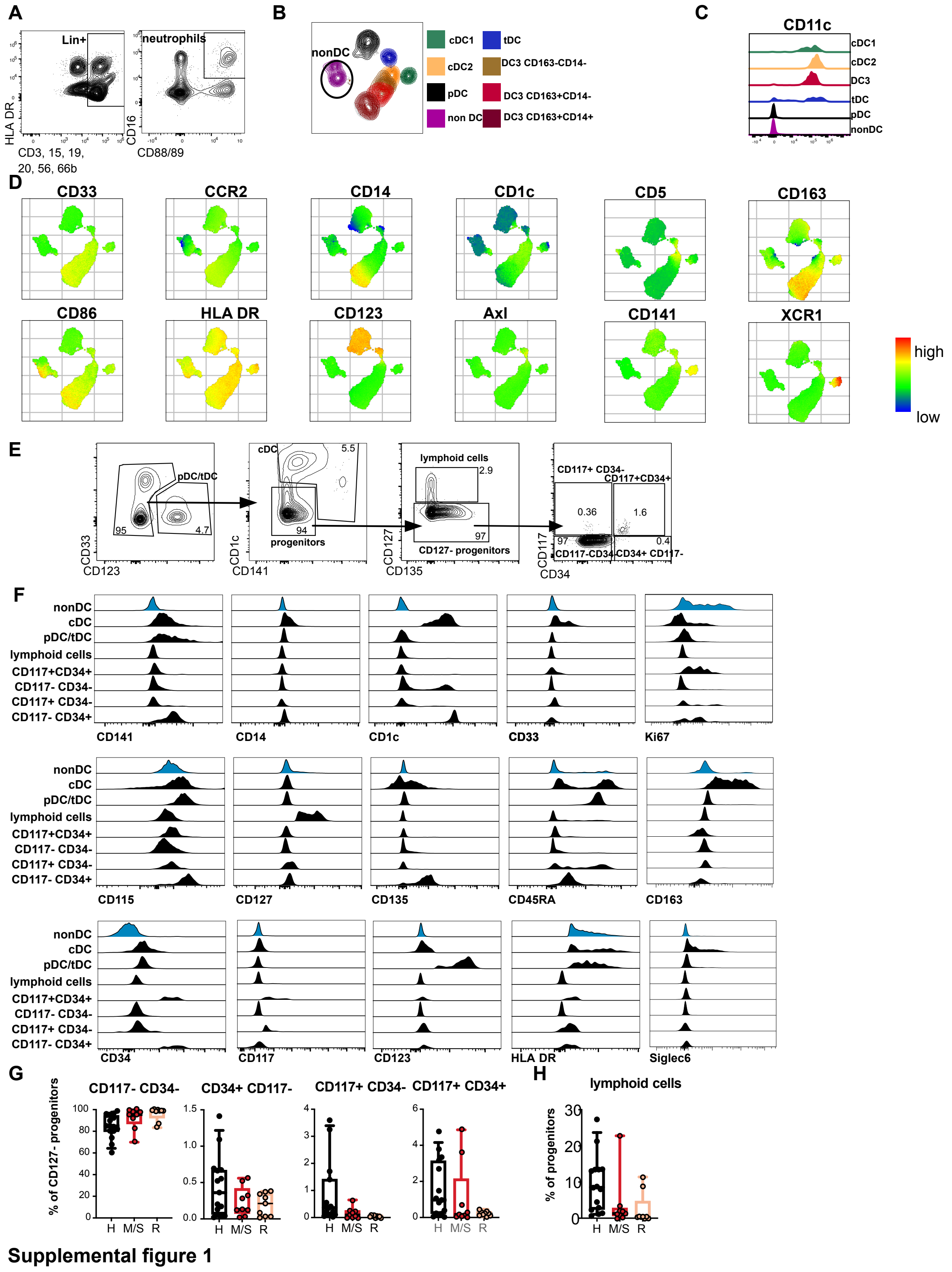


**Fig. S1. Characterization of HLADR+ non-DC population expanded in COVID-19 patients**

(A) Gating strategy for neutrophils in the blood: Within the lineage (CD3, CD15, CD19, CD20, CD56, CD66b) positive cells neutrophils were gated as $CD16^{+}$ and $CD88/89^{+}$.

(B) UMAP clustering of one representative COVID-19 patient. Overlay of gated cDC1 (green, $CD141^{+}$), cDC2 (orange, $CD1c^{+}$, $CD5^{+}$), DC3 (brown, red, dark red, $CD1c^{+}$, $CD5^{–}$, $CD163^{+/–}$, $CD14^{+/–}$), pDC (black, $CD123^{+}$), tDC (blue, $CD123^{+}$, $Siglec1^{+}$, $Axl^{+}$) and non-DC (purple, $HLADR^{+}$, Lin-, $CD141^{–}$, $CD1c^{–}$) populations. (C) Representative histograms of CD11c expression in cDC1, cDC2, DC3, tDC, pDC and non-DC in a patient with moderate COVID-19. (D) Expression of several surface markers overlayed in the UMAP from (B). Shown is the expression of CD33, CCR2, CD14, CD1c, CD5, CD163, CD86, HLADR, CD123, Axl, CD141, XCR1 indicated by colour scale (red = high expression, green = intermediate, blue = low expression). (E) Gating strategy for identification of progenitor populations in the blood. Cells are pregated on Lin- (CD3, CD15, CD19, CD20, CD56, CD66b, CD88, CD89), $HLADR^{+}$ living cells. pDCs and tDCs are excluded via gating on $CD123^{–}$ cells followed by exclusion of cDCs by gating on $CD1c^{–}$, $CD141^{–}$ cells. These progenitors are then separated into lymphoid cells ($CD127^{+}$) and $CD127^{–}$ progenitors. Here, cells can be differentiated by their expression of CD117 and CD34 into four quadrants: $CD117^{+}$ $CD34^{–}$, $CD117^{+}$ $CD34^{+}$, $CD117^{–}$ $CD34^{+}$, $CD117^{–}$ $CD34^{–}$. (F) Representative histograms of marker expression of non-DCs, cDCs, pDC/tDCs, lymphoid cells, $CD117^{+}$ $CD34^{+}$, $CD117^{–}$ $CD34^{–}$, $CD117^{+}$ $CD34^{–}$, $CD117^{–}$ $CD34^{+}$ progenitor cells, gated according to (E) of one COVID-19 patient. Expression of CD141, CD14, CD1c, CD33, Ki67, CD115, CD127, CD135, CD45RA, CD163, CD34, CD117, CD123, HLADR and Siglec6 is shown. (G) Percentage of $CD117^{–}$ $CD34^{–}$, $CD34^{+}$ $CD117^{–}$, $CD117^{+}$ $CD34^{–}$, $CD117^{+}$ $CD34^{+}$ cells of total $CD127^{–}$ progenitor cells. (H) Percentage of $CD127^{+}$ lymphoid cells of the progenitor cells are shown. Healthy donors (=H, black symbols, n=12), mild/moderate and severe COVID-19 patients (=M/S, red, n=9) and recovered patients (=R, orange, n=9).

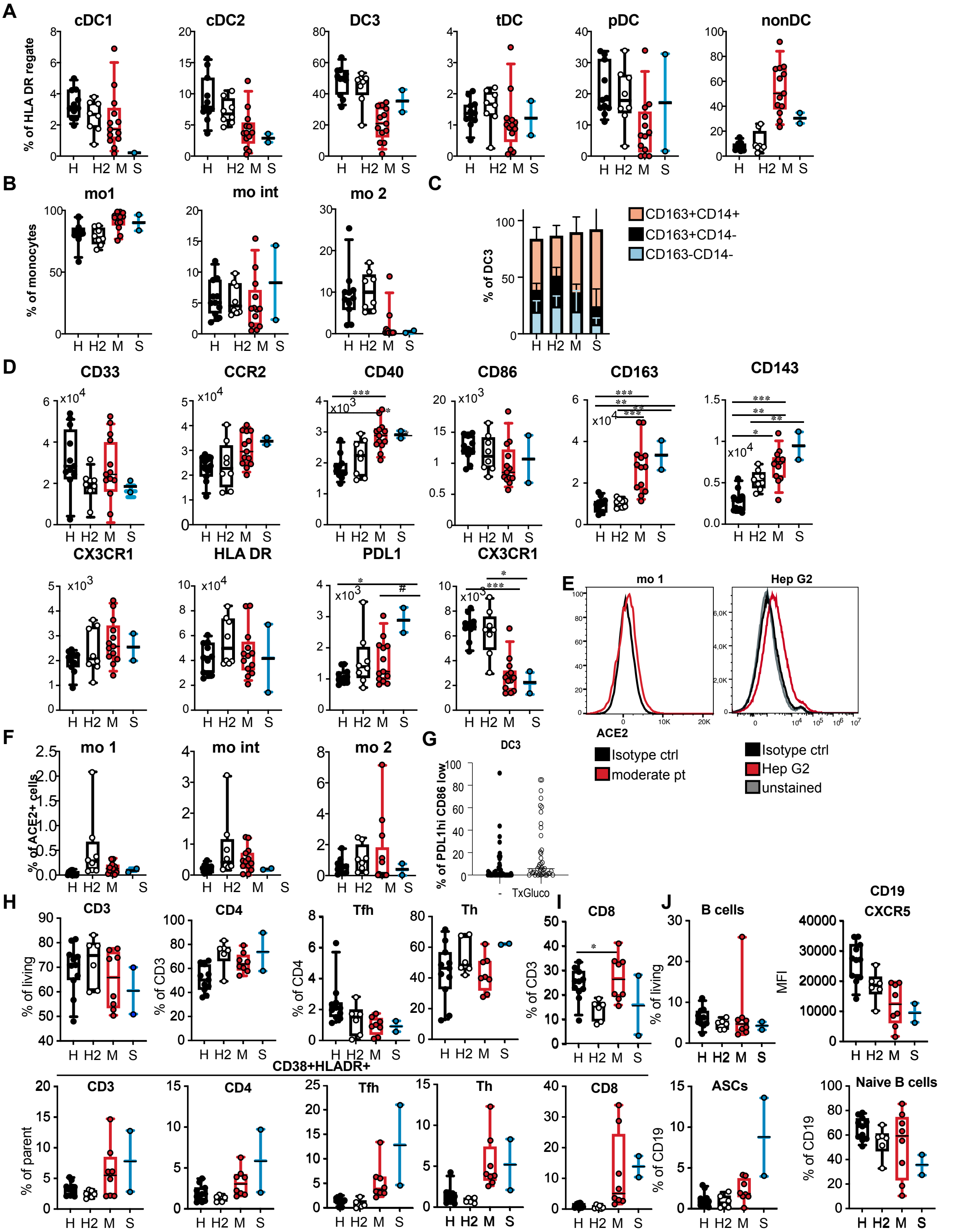


**Supplemental figure 2**

**Fig. S2. Frequency and phenotype of DC, monocyte, T cell and B cell subpopulations in young vs. old healthy donors and COVID-19 patients**

(A) Relative frequencies of DC subsets and non-DCs within the DC gate are shown.

(B) Relative frequencies of classical monocytes (mo 1), intermediate monocytes (mo int) and non-classical monocytes (mo 2) within the monocyte gate are shown. (C) Relative frequencies of DC3 subtypes identified by CD163 and CD14 expression are shown (mean and SD). (D) Surface expression (MFI) of several markers shown in mo 1. (E) Representative histogram of ACE2 expression in mo 1 in a moderate COVID-19 patient (red) and the isotype control (black) and of ACE2 expression in Hep G2 cells (red), unstained Hep G2 cells (grey) and the isotype control (black). (F) Relative frequencies of ACE2 positive cells in mo 1, mo int and mo 2. (A-F) Healthy patients (=H, black, n=11), aged healthy patients (=H2, white, n=8), mild/moderate COVID-19 pts (=M, red, n=13) and severe COVID-19 patients (=S, blue, n=2). (G) Frequencies of PD-L1$^{hi}$ CD86$^{lo}$ DC3 in patients receiving glucocorticoid therapy (white) and not receiving glucocorticoid therapy (black) are shown (n=86). (H, I) Frequencies of the indicated T cell populations. (J) Frequencies of the indicated B cell populations and CXCR5 MFI values. (H-J) Healthy patients (=H1, black, n=11), aged healthy patients (=H2, white, n=6), mild/moderate COVID-19 pts (=M, red, n=8) and severe COVID-19 patients (=S, blue, n=2). Kruskal-Wallis Test with Dunn's correction, or ANOVA and Tukey's test was used, *p<0.05, **p<0.01, ***p<0.001.

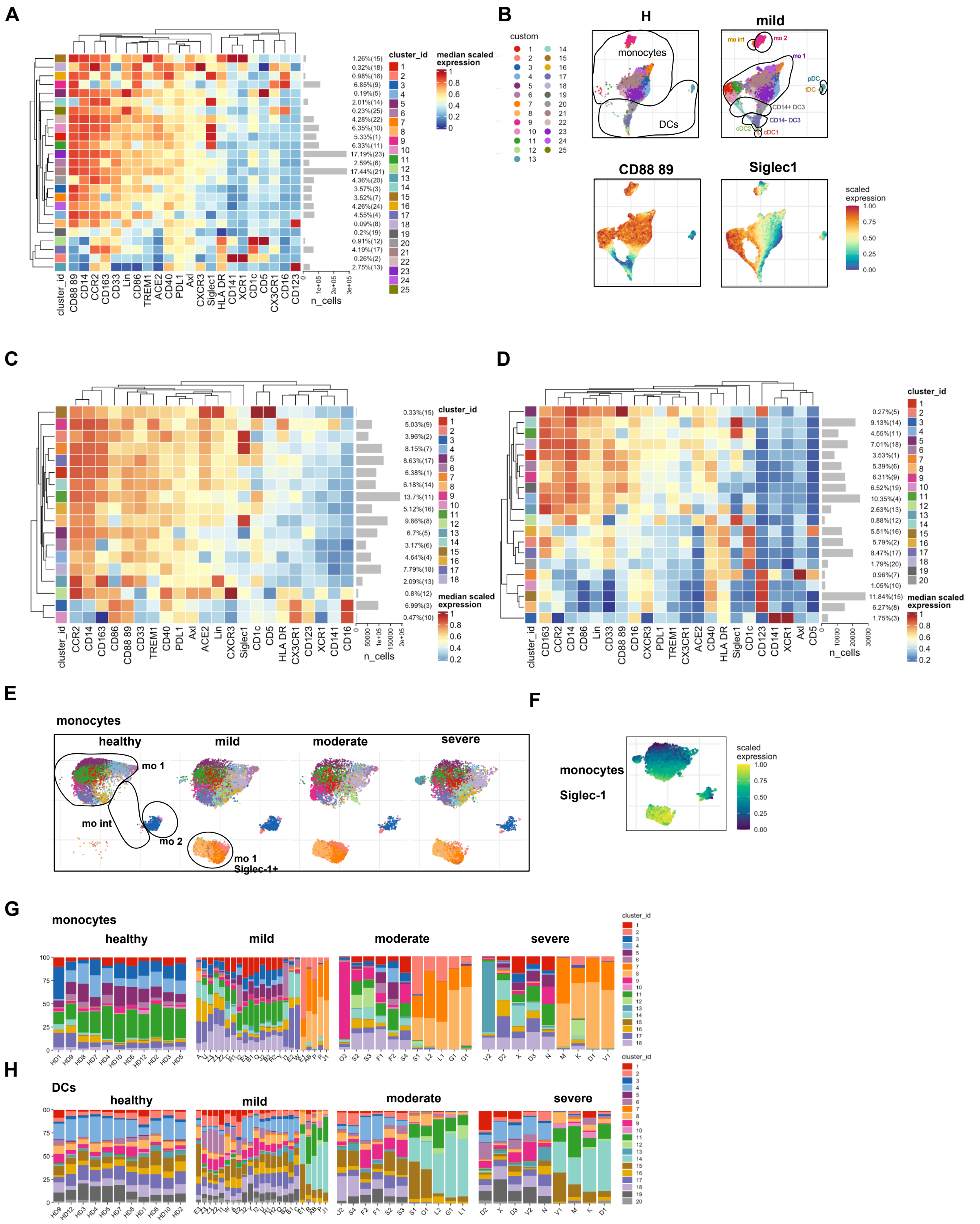


**Supplemental figure 3**

**Fig. S3. Analysis of DC and monocyte subpopulations by phenograph clustering**

(A) Heatmap of marker expression in phenograph clusters and (B) UMAP of HLADR$^+$/intermediate Lin$^-$ cells reclustered after exclusion of all undefined cells. UMAPs of pooled data from healthy controls and COVID-19 patients with mild disease are shown with annotation of monocyte and DC subpopulations. Color overlays indicate scaled marker expression. (C) Heatmap of marker expression in phenograph clusters of reclustered reclusterednd (D) of reclustered reclusteredAPs of reclustered reclusteredith Phenograph clusters indicated by colors are shown separately for the indicated groups with annotation of monocyte subpopulationColor Colour overlay indicates scaled Siglec-1 expression in COVID-19 patients with moderate disease. (G and I) Frequencies of phenograph clusters in individual patients grouped by disease severity and controls. Letter indicates patients, numbers indicate consecutive sampling timepoints. Healthy donors were numbered.

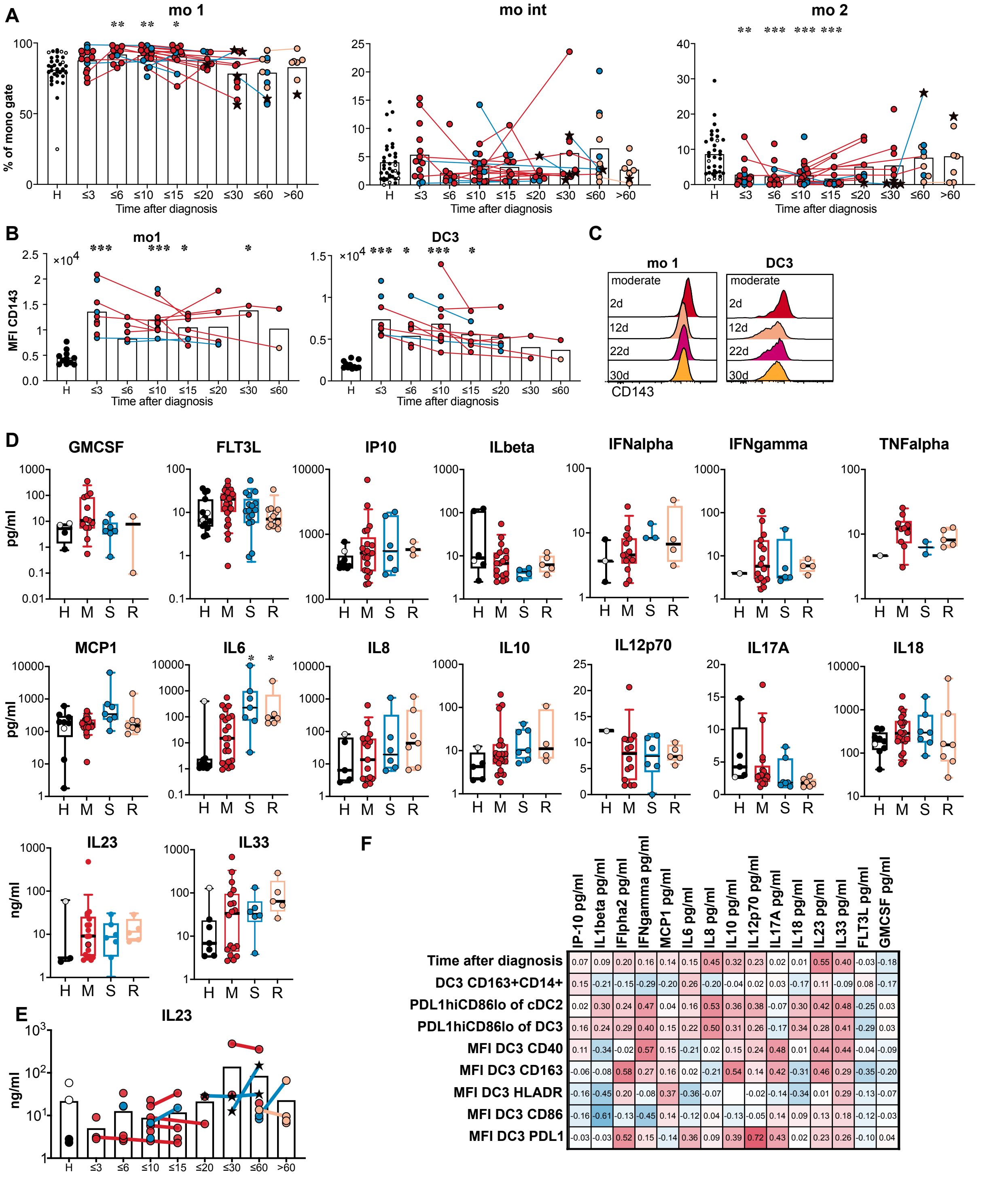


**Supplemental figure 4**

**Fig. S4. Monocyte subsets and plasma cytokines in COVID-19 patients compared to controls**

(A) Frequency of mo 1, mo int and mo 2 of all monocytes at different timepoints after diagnosis. (B) CD143 expression at different timepoints after diagnosis in mo 1 and

DC3. (A, B) Connected lines represent multiple measurements of the same donor at different time points. Columns indicate the mean. Kruskal-Wallis test with Dunn's correction. *p<0.05, **p<0.01, ***p<0.001, ****p<0.0001. (C) Representative histograms of CD143 expression in mo 1 and DC3 in a patient with moderate COVID-19 at the indicated time points after diagnosis. (D) Plasma concentrations (pg/ml) of plasma cytokines in healthy patients (=H, black, n=1-15), mild/moderate COVID-19 pts (=M, red, n=10-30), severe COVID-19 pts (=S, blue, n=2-17) and recovered (=R, orange, n=5-13) measured at the first timepoint after diagnosis. (E) Plasma concentrations (pg/ml) of IL-23 at different grouped time points after diagnosis. Connected lines represent multiple measurements of the same donor at different time points. Columns indicate the mean. Kruskal-Wallis test with Dunn's correction, n=44. *p<0.05, **p<0.01, ***p<0.001. (F) Spearman correlation of plasma concentrations of cytokines at all timepoints with innate parameters at the same sampling time points (n=9-94).

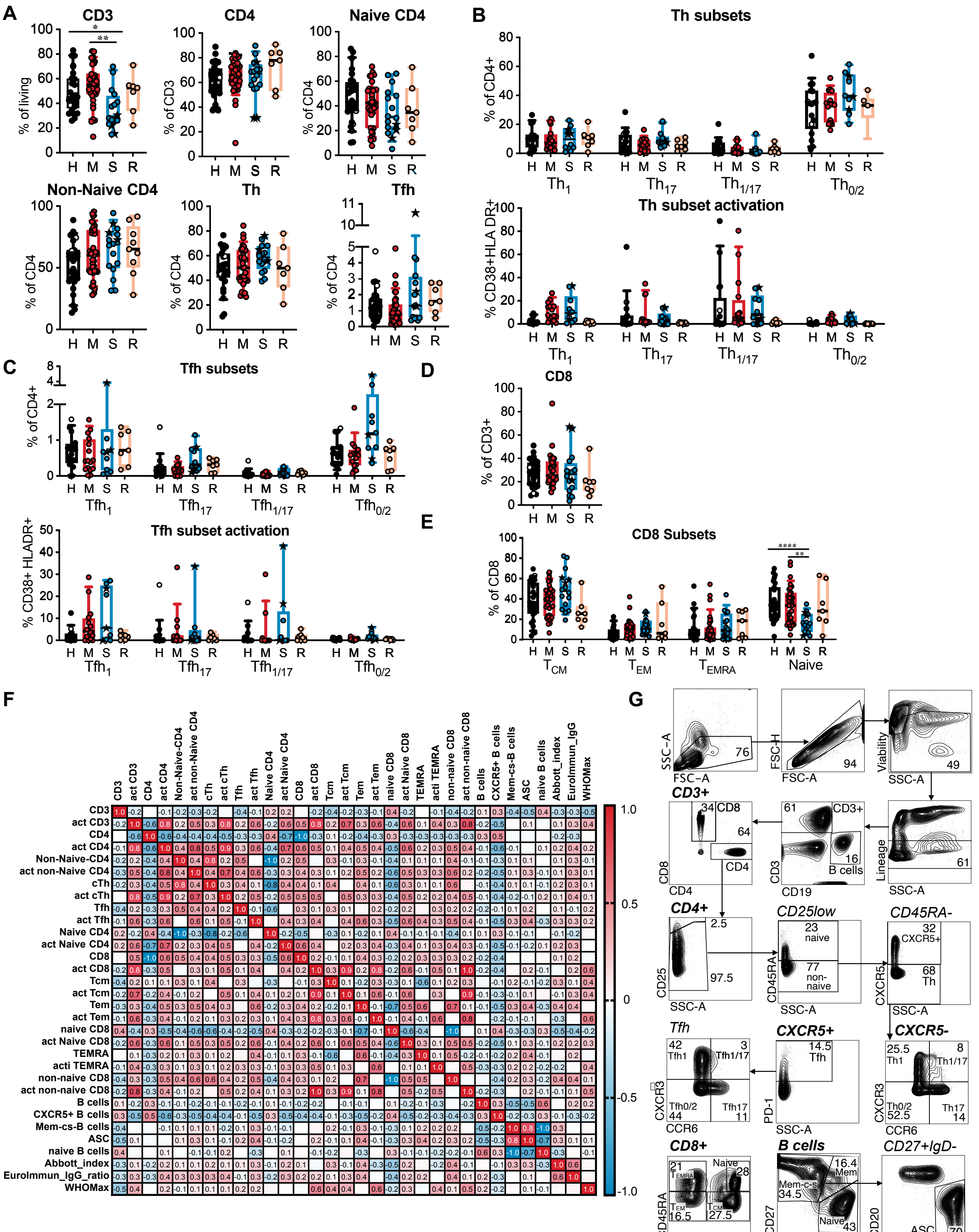


Supplemental figure 5

**Fig. S5. Frequency and activation of T cell subpopulations in COVID-19 patients compared to controls**

(A) Frequencies of the indicated T cell populations in healthy/non-COVID controls (H, n=24) and acute COVID-19 patients with mild/moderate (M, n=35) or severe disease

(S, n=16) at the first analysis timepoint and recovered patients (R, n=7). (B) Frequencies of Th cell subsets (Th1: $CXCR3^+$ $CCR6^-$, Th17: $CXCR3^-$ $CCR6^+$, Th1/17: $CXCR3^+$ $CCR6^+$, Th0/2: $CXCR3^-$ $CCR6^-$) in $CD4^+$ T cells. Percentage of $CD38^+$ $HLADR^+$ activated cells within the indicated Th cell subsets. (C). Frequencies of Tfh cell subsets in $CD4^+$ T cells. Percentage of $CD38^+$ $HLADR^+$ activated cells within the indicated Tfh-like cell subsets. (B, C) H, n=22; M, n=16; S, n=10; R, n=7. (D) Percentage of $CD8^+$ T cells. (E) Frequencies of $CD8^+$ naïve and memory subsets with $CD8^+$ T cells. (D, E) H, n=24; M, n=35; S, n=16; R, n=7. Kruskal-Wallis test with Dunns correction, *$p<0.05$, **$p<0.01$, ***$p<0.001$. (F) Spearman correlation of adaptive parameters at time after diagnosis 10 to 25 days (n=35-42). (G) Gating strategy for T and B cell subpopulations shown for one COVID-19 patient.
